# Timing of organ initiation is crucial for robust organ size

**DOI:** 10.1101/2020.03.08.982629

**Authors:** Mingyuan Zhu, Weiwei Chen, Vincent Mirabet, Lilan Hong, Simone Bovio, Soeren Strauss, Erich M. Schwarz, Satoru Tsugawa, Zhou Wang, Richard S. Smith, Chun-Biu Li, Olivier Hamant, Arezki Boudaoud, Adrienne H. K. Roeder

**Affiliations:** Weill Institute for Cell and Molecular Biology and School of Integrative Plant Science, Section of Plant Biology, Cornell University, Ithaca NY 14853, USA; Laboratoire de Reproduction et Développement des Plantes, Université de Lyon, UCB Lyon 1, ENS de Lyon, INRA, CNRS, 46 Allée d’Italie, 69364 Lyon Cedex 07, France; Department of Comparative Development and Genetics, Max Planck Institute for Plant Breeding Research, Cologne, 50829 Germany; Department of Molecular Biology and Genetics, Cornell University, Ithaca, NY 14853, USA; Department of Mathematics, Stockholm University, 106 91 Stockholm, Sweden; Key Laboratory of Horticulture Science for Southern Mountains Regions of Ministry of Education; College of Horticulture and Landscape Architecture; Southwest University, Beibei, Chongqing 400715, China; Academy of Agricultural Sciences of Southwest University; State Cultivation Base of Crop Stress Biology for Southern Mountainous Land of Southwest University, Beibei, Chongqing 400715, China; Institute of Nuclear Agricultural Sciences, College of Agriculture & Biotechnology, Zhejiang University, Hangzhou, Zhejiang 310058, China; Graduate School of Biological Sciences, Nara Institute of Science and Technology, Ikoma Nara 630-0192, Japan

## Abstract

Organs precisely regulate their size and shape to ensure proper function^1–6^. The contribution of organ initiation timing to final organ size and shape is often masked by compensatory adjustments to growth later in development^7–9^. Here we show that DEVELOPMENT RELATED MYB-LIKE1 (DRMY1) is required for both proper organ initiation timing and growth leading to robust sepal size in *Arabidopsis.* Within each *drmy1* flower, the initiation of some sepals is variably delayed. Late-initiating sepals in *drmy1* mutants remain smaller throughout development resulting in variability in sepal size. DRMY1 focuses the spatiotemporal signaling patterns of the plant hormones auxin and cytokinin, which jointly control the timing of sepal initiation. Contrary to expectation, our findings demonstrate that timing of organ initiation contributes to robust organ size throughout development.

## Main Text

Development is remarkably reproducible, generally producing the same organ with invariant size, shape, structure, and function in each individual. For example, the two arms of a person match in length with an accuracy of 0.2% ^1^, mouse brains vary in size by only about 5% ^2^, and *Arabidopsis* floral organs are strikingly uniform ^3^. Defects in organ size control mechanisms contribute to many human diseases including hypertrophy and cancer ^4, 5^. Uniformity of fruit size is an important criterion for packaging and shipping fruit to market ^6^. In this context, robustness is the ability to form organs reproducibly despite perturbations, such as stochasticity at the molecular and cellular level as well as environmental fluctuations ^10^. Robustness has fascinated biologists since Waddington brought the issue to prominence in 1942 ^11^. One proposed scenario for achieving organ size robustness is that organs can sense their size and compensate through adjusting their growth rates or maturation time until the organ has reached the correct size ^7^. For example, in *Drosophila*, damaged or abnormally growing imaginal disks activate the expression of *Drosophila* insulin-like peptide8 (DILP8), which delays metamorphosis and thus allows damaged disks to reach the correct size ^8, 9^. These compensatory mechanisms can mask early-stage defects, which raises the question of whether events early in organogenesis play any role in ensuring organ size robustness.

Sepals, the outermost floral organs, are a good model system for investigating the mechanisms of organ size robustness because individual plants can produce more than 50 invariant flowers. This allows statistical assessment of organ size uniformity within a single organism that cannot be achieved in most model systems. Sepals arise from floral meristems (FM, stem cells that give rise to floral organs), which initiate from the periphery of the inflorescence meristem (IM, stem cells at the tip of the plant that give rise to the flowers; Fig. 1A). On the flank of *Arabidopsis thaliana* floral meristems, four sepals initiate and rapidly grow to cover that flower. The four growing sepals in a flower must maintain the same size and shape to enclose and protect developing reproductive organs throughout growth before the flower blooms ^12^; thus, continuous robustness of size and shape is required for sepal function (Fig. 1A-B). We established a nomenclature for the four sepals in a flower. The sepal closest to the IM is the inner sepal, while the sepal opposite, farthest from the IM, is the outer sepal. The two sepals on the sides are lateral sepals (Fig. 1A).

**Fig. 1.**
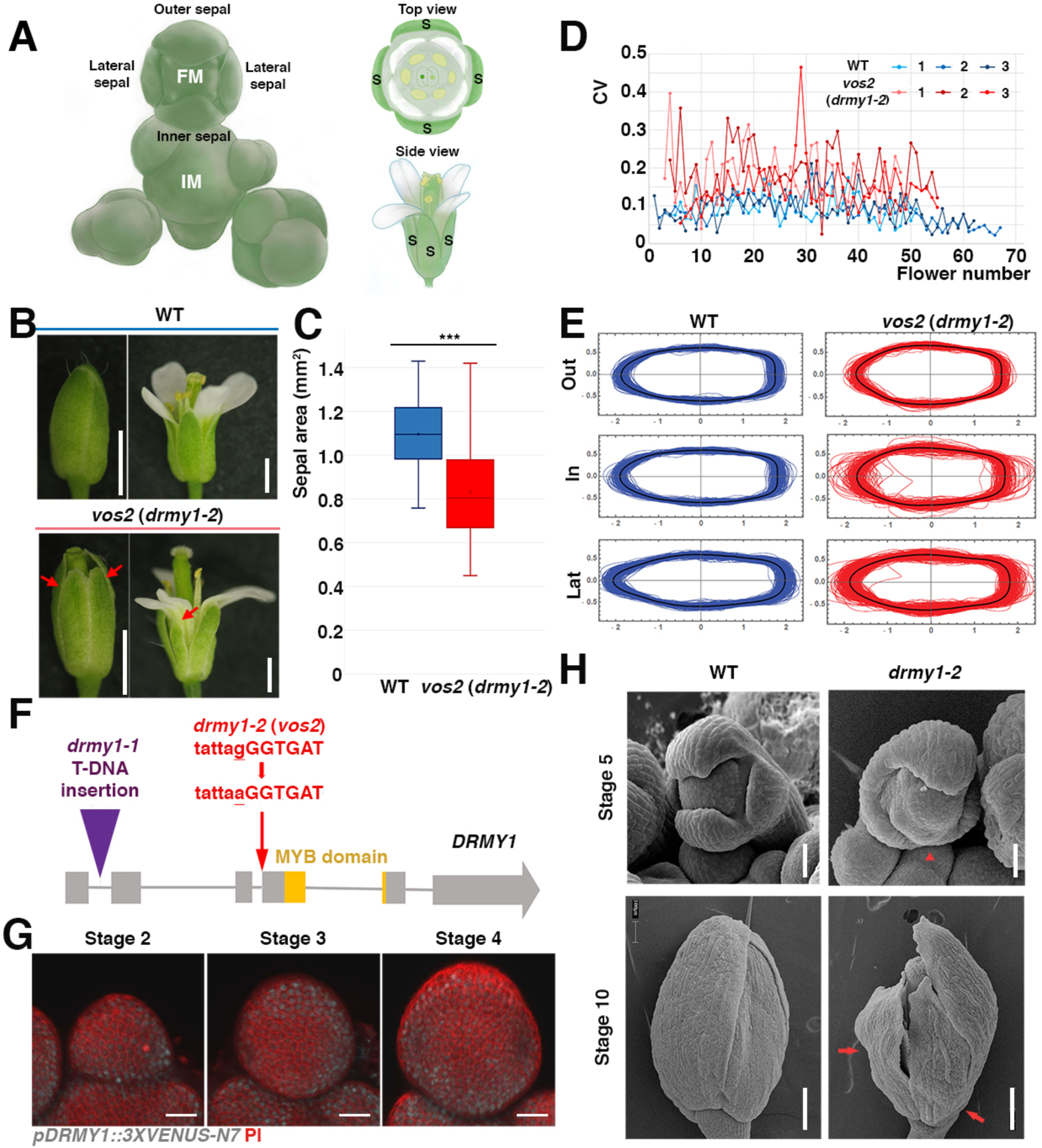
Mutations in the *DRMY1* gene lead to increased variation in sepal size and shape. (A) Anatomical diagram. Floral meristems (FM) emerge from the inflorescence meristem (IM). Four sepals (S) initiate evenly from the periphery of the floral meristem. The inner sepal is closest to the IM, the outer sepal farthest from the IM, and the lateral sepals on the sides. Note that throughout development, the sizes of sepals are always similar. (B) Wild-type (WT) and *vos2* (*drmy1-2*) flowers as closed buds (stage 12, left) and after blooming (stage 15, right). Red arrow: smaller sepals within each *vos2* (*drmy1-2*) flower. Scale bars:1 mm. (C) The sepal area distribution is wider for the *vos2* (*drmy1-2*) mutant compared to WT. Area variability was quantified by the coefficient of quartile variation (CQV) and was significantly higher in *vos2* (Permutation test ***: p-value < 0.001). The average area of *vos2* sepals was significantly lower than wild type (Permutation test ***: p-value < 0.001). The boxes extend from the lower to upper quartile values of the data, with a line at the median. And the whiskers extend past 1.5 of the interquartile range. n = 100 for both WT and *vos2* (*drmy1-2*) 10^th^ to 25^th^ flowers along the main branch. Outer, inner, and lateral sepals were pooled together. (D) Coefficient of variation (CV) calculated for the areas of the four sepals in each single flower. Sequential flowers along the main branch of the stem (flower number on the x-axis) were measured at stage 14. Three replicates are included for both WT (blue) and *vos2* (*drmy1-2*, red) mutants. (E) *vos2* (*drmy1-2*) mutants also exhibit higher variation in inner and lateral sepal shape. Superimposed outlines of stage 14 sepals from WT and *vos2* (*drmy1-2*) mutants were normalized by sepal area. The black outline is the median. (F) Gene model for the *DRMY1* gene (AT1G58220). Orange box indicates the location of the MYB domain. A G to A point mutation was identified at an exon and intron junction in the *vos2* (*drmy1-2*) mutant. *drmy1-1* was reported with a T-DNA insertion in the first intron. (G) Expression pattern of *pDRMY1::3XVENUS-N7* (white). The expression of nuclear localized VENUS driven by the *DRMY1* promoter was observed in young stage flowers, especially the peripheral zones. Cell walls were stained with propidium iodide (PI, red). Scale bar: 20 µm. (H) Scanning electron micrographs show that the sepal size variability phenotype can be observed at early stages (stage 5) and remains visible through late stages. Red arrowhead: delayed sepal initiation at stage 5; Red arrows: smaller sepals within the single *drmy1-2* flowers; Scale bar at top panel: 30µm; Scale bar at bottom panel: 200 µm.

### Mutations in *DRMY1* cause variability in sepal size

Robustness mechanisms can be identified by screening for mutants with increased variability ^13, 14^. Accordingly, we screened for mutants exhibiting variable sizes or shapes of the sepals, thus disrupting robustness ^14^. Our previous analysis of the *<u>v</u>ariable <u>o</u>rgan <u>s</u>ize and shape1 (vos1*) mutant revealed that highly variable cell growth is averaged in time and space to create robust organs ^14^. From that mutant screen, we also isolated the *<u>v</u>ariable <u>o</u>rgan <u>s</u>ize and shape2 (vos2*) mutant which had sepals of different sizes within the same flower. Consequently, *vos2* sepals failed to form a complete barrier to protect the inner reproductive organs (Fig. 1B). To exclude the possibility that the variability arose from the altered sepal number, we counted the number of sepals produced in *vos2* flowers and found that it was largely unaffected, with 4 sepals present in >92% (164/177) of *vos2* flowers (compared to 100% (207/207) of wild-type flowers; Fig. S1B). Next, we quantified the size distribution of mature sepals from many mutant plants, in flowers with 4 sepals. We found that *vos2* mutant sepals had increased variability in area and reduced average area compared to wild type (Fig. 1C, S1C and S1D). We further assayed individual flowers developing sequentially along the main branch of a single plant and found that the area of the four sepals in each *vos2* flower consistently exhibited higher coefficients of variation (CV) and smaller averages in sepal area than wild type (Fig. 1D and Fig. S1A). Additionally, the shapes of *vos2* inner and lateral mutant sepals were more variable than wild-type sepals, after normalizing for size (Fig. 1E and Fig. S1E). VOS2 promotes sepal growth as its disruption led to smaller sepals. One might hypothesize that higher variance originates from altered average size. However, we have previously shown that decreasing sepal sizes does not automatically lead to increased sepal size variability in mutants ^14^. Moreover, for both wild type and *vos2* mutants, we saw no correlation between average and CV in sepal size (Fig. S1C and S1D). Thus, the loss of regularity is not simply a side effect of decreased average sepal size. VOS2 itself is required for robustness of sepal size and shape.

Map-based cloning of *vos2* identified a G to A point mutation in a splice acceptor site of the gene (AT1G58220) encoding the MYB domain protein <u>D</u>EVELOPMENT <u>R</u>ELATED <u>MY</u>B LIKE 1 (DRMY1; Fig. 1F). This point mutation caused altered splicing resulting in premature stop codons (Fig. S1F), as well as a dramatic decrease of *DRMY1* transcript level (Fig. S1G). A T-DNA insertion allele, *drmy1-1*, was recently reported as broadly affecting cell expansion ^15^. Thus, we renamed *vos2* as *drmy1-2*. To verify that mutations in *DRMY1* caused the variable sepal size and shape phenotype, we observed that the *drmy1-1* T-DNA insertion allele also exhibited sepal size variability, and that *drmy1-1* and *drmy1-2* alleles failed to complement, indicating they were alleles of the same gene (Fig. S1H). Furthermore, expression of *DRMY1* under its endogenous promoter rescued the sepal variability phenotypes (20/21 rescued in T1), confirming the role of *DRMY1* in regulating sepal robustness (Fig. S1H).

To determine when DRMY1 functions in sepal robustness, we examined reporters for DRMY1 expression. The DRMY1-mCitrine fusion protein (*pDRMY1::DRMY1-mCitrine*) rescued the *drmy1-2* mutant phenotypes (21/23 rescued in T1), indicating that the fusion protein is functional (Fig. S1H). The DRMY1 reporters were expressed broadly within young flowers, floral meristems and inflorescence meristems (Fig. 1G and S1I). DRMY1 reporters had somewhat higher expression within the periphery of developing floral and inflorescence meristems, hinting that DRMY1 might function in organ initiation.

### Sepal primordium initiation is variably delayed in *drmy1-2* mutants

Since *DRMY1* reporters were expressed before and during sepal primordium initiation, we used scanning electron microscopy to determine the stage at which the defect in sepal size robustness was first visible in *drmy1* mutants. Sepal variability in *drmy1-2* arose during initiation and was visible throughout flower development (Fig. 1H). In wild type, the four sepals were the first organ primordia to initiate at the periphery of the floral meristem. At the same stage in *drmy1-2*, the flowers exhibited a normal-looking outer sepal. However, the inner and lateral sepal primordia were often absent or appeared smaller than wild type, suggesting that their initiation was delayed (Fig. 1H). As mentioned above, >92% of *drmy1-2* flowers had four sepals at maturity; this was consistent with a delay rather than a block of sepal initiation.

To determine how much the timing of sepal initiation is actually delayed in the *drmy1-2* mutant, we live imaged wild-type and the *drmy1-2* mutant flowers throughout the initiation of sepal primordia (Fig. 2A and B, and Movie S1 and S2). We defined the bulging of sepal primordia out from the floral meristem as the morphological initiation event, which we detected by observing the Gaussian curvature of the meristem surface. A clear band of positive curvature (red in the heat map) at the flank of the floral meristem indicated the initiation of the sepal (Fig. 2C and D). The initiation of the outer sepal occurred first, and was set as the starting point, followed by the inner and then lateral sepals. For wild type, the time intervals between the initiation of outer and inner sepals were always around 6 hours (Fig. 2 A, C, and E). Within 12 hours after the initiation of the outer sepal, the two lateral sepals initiated (Fig. 2A, C, and F). The sepal primordia then rapidly grew to cover the floral meristem by 30 hours (Fig. 2A).

**Fig. 2.**
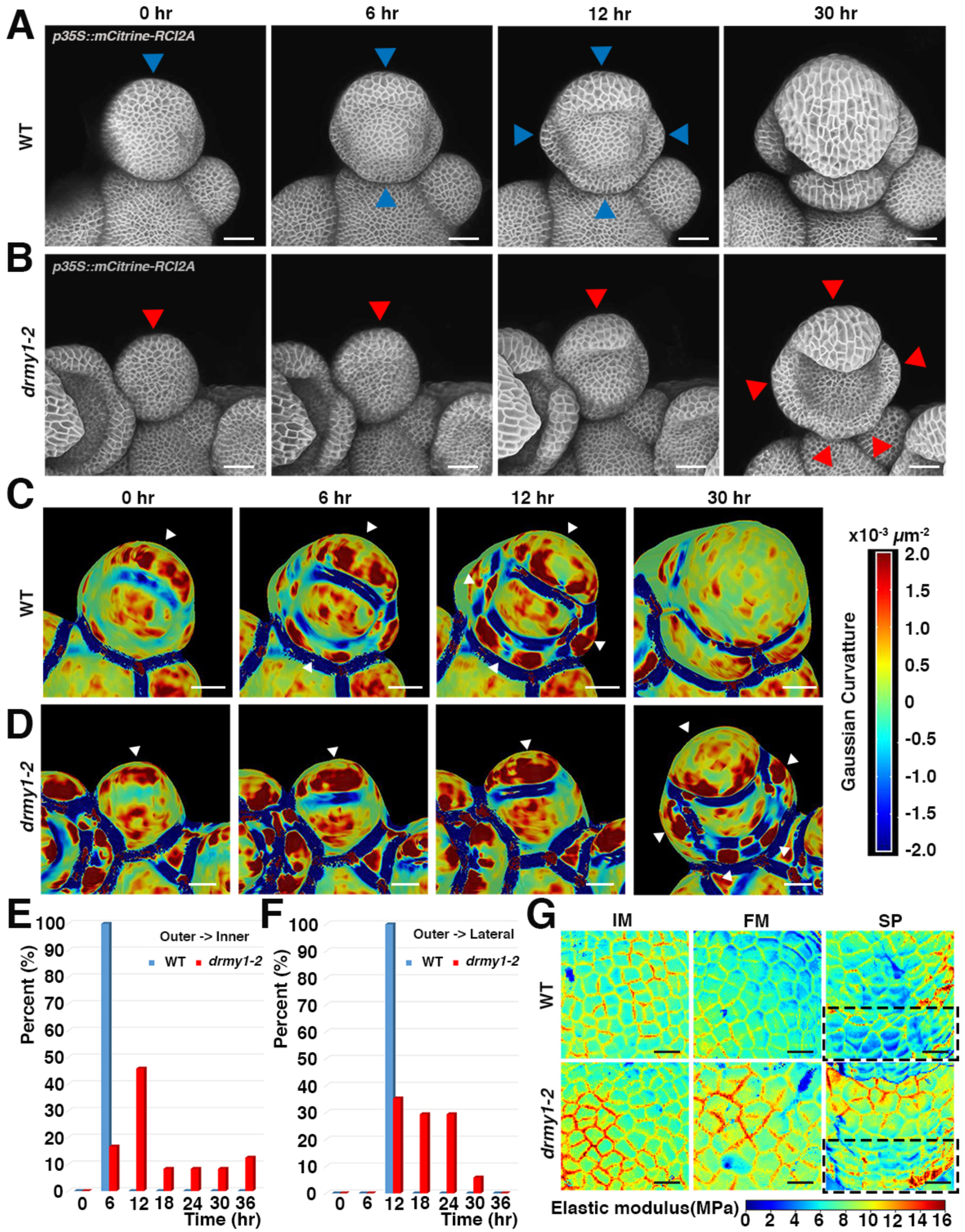
Sepal initiation is variably delayed in the *drmy1-2* mutant. (A and B) Live imaging of sepal initiation in wild type (A) and *drmy1-2* (B). Plasma membrane marker (*p35S::mCitrine-RCI2A*) is shown in greyscale. The small bulge-out is defined as sepal initiation. Blue arrowheads in WT and red arrowheads *drmy1-2* indicate initiated sepals. Scale bars: 25 µm. (C and D) Gaussian curvature heatmap detecting changes in curvature associated with initiation for the live imaging sequences shown in A and B. The red color represents the dome shape while the blue color represents the saddle shape. Thus, a strong red band at the periphery (white arrowheads) reveals initiated sepals. Scale bars: 25 µm. (E) Histogram showing the time interval between the initiation of the outer sepal and the inner sepal in each flower. n = 12 for WT flowers and 12 for *drmy1-2* flowers. Note WT inner sepals initiate robustly 6 hours after the outer sepals, while the time is variable and generally longer in *drmy1-2*. (F) Histogram showing the time interval between the initiation of the outer sepal and the lateral sepals for each flower. (G) Atomic Force Microscopy measurement of cell wall stiffness for centers of inflorescence meristems (IM), centers of floral meristems (FM), and peripheries of floral meristems where sepal primordia emerge (SP; highlighted with black dotted boxs) of WT and the *drmy1-2* mutants. In the heatmap of apparent elastic modulus shown here, red indicates stiffer and blue indicates softer. (n = 11) Scale bars: 10 µm.

In contrast, for the *drmy1-2* mutant, the time intervals for the initiation of inner and lateral sepals were elongated and more variable (Fig. 2B, D, E and F). In *drmy1-2*, the inner sepals initiated anywhere from 6 to 36 hours after the outer sepals (Fig. 2E). Likewise, the lateral *drmy1-2* sepals initiated from 12 to 30 hours after the outer sepal (Fig. 2F). Initiation of the lateral sepals occasionally occurred before the initiation of the inner sepal in *drmy1-2* flowers. Frequently, two sepal primordia appeared to form instead of one at the inner position of *drmy1-2* flowers (e.g. highlighted with red arrowheads in Fig. 2B). However, further live imaging revealed that most of these fused to form a single sepal with two tips, resulting in the four sepals finally observed in *drmy1-2* flowers (Fig. S1J). In addition, it took much more time for *drmy1-2* sepal primordia to cover the whole floral meristem (Movie S2). We performed our analysis relative to the initiation of the outer sepal, which appeared normal.

### Stiffer cell walls in *drmy1-2* mutants correlate with delayed sepal initiation

*DRMY1* encodes a MYB domain protein, and most MYB domain proteins function as transcription factors. To identify the biological processes that are regulated by DRMY1 to promote robust timing of sepal primordium initiation, we performed an RNA-seq experiment comparing inflorescences and flowers of *drmy1-2* mutants to wild type. Gene ontology (GO) term analysis of the differentially expressed genes revealed an enrichment of biological processes including “cell wall modification”, “response to hormone stimulus” and “cellular metabolic process” (Fig. S2D and E; Dataset S1). Plant cell walls become softer through cell wall modification during primordium initiation to allow outgrowth. Genetically stiffening the cell wall is sufficient to block the initiation of organ primordia ^16, 17^. Our RNA-seq data suggested that cell wall stiffness might be changed in *drmy1-2* mutants due to altered cell wall modifications. To determine whether the delayed organ initiation in *drmy1-2* might result from a stiffer cell wall, we first used <u>A</u>tomic <u>F</u>orce <u>M</u>icroscopy (AFM) to quantify the cell wall stiffness of the sepal primordia, the floral meristem and the inflorescence meristem. For all three, the cell wall was significantly stiffer in the *drmy1-2* mutant (higher average apparent elastic modulus; Fig. 2G, Fig. S2A), consistent with the delay of primordium initiation. We also used osmotic treatment to assess cell wall stiffness by quantifying the shrinkage of cell walls when internal turgor pressure was decreased. Osmotic treatment of wild-type and *drmy1-2* developing sepals further confirmed that cell walls were stiffer in the *drmy1-2* mutant (Fig. S2B and C). Our data is consistent with the model that stiffer cell walls in *drmy1-2* led to delayed sepal initiation, and consequently higher sepal size variability.

### Variably delayed initiation disrupts sepal size robustness throughout development

To test whether delayed sepal initiation decreases sepal size throughout flower development, we live imaged sepals from their initiation throughout their development over 11 days (Fig. 3A-B and Fig. S3E). In wild type, after robust initiation, the sepals maintained equivalent sizes so that flowers remained closed throughout development (Fig. 3A). At the end of our live imaging series, sepal sizes were equivalent (Fig. S3E). In contrast, in *drmy1-2* flowers, sepals with delayed initiation remained smaller than other sepals throughout development, so that flowers remained open throughout development (Fig. 3B). At the end of our live imaging series, these sepals had variable sizes (Fig. S3E). Validating this result, our previous quantification of mature sepal size showed that in general, the *drmy1-2* inner and lateral sepals had a more severe decrease in size relative to wild type than outer sepals, correlating with their delayed initiation (Fig. S3F and S3G). These results indicate that precisely timed initiation is critical for robustness in organ size and raises the question of why compensation is inefficient in *drmy1* mutants.

**Fig. 3.**
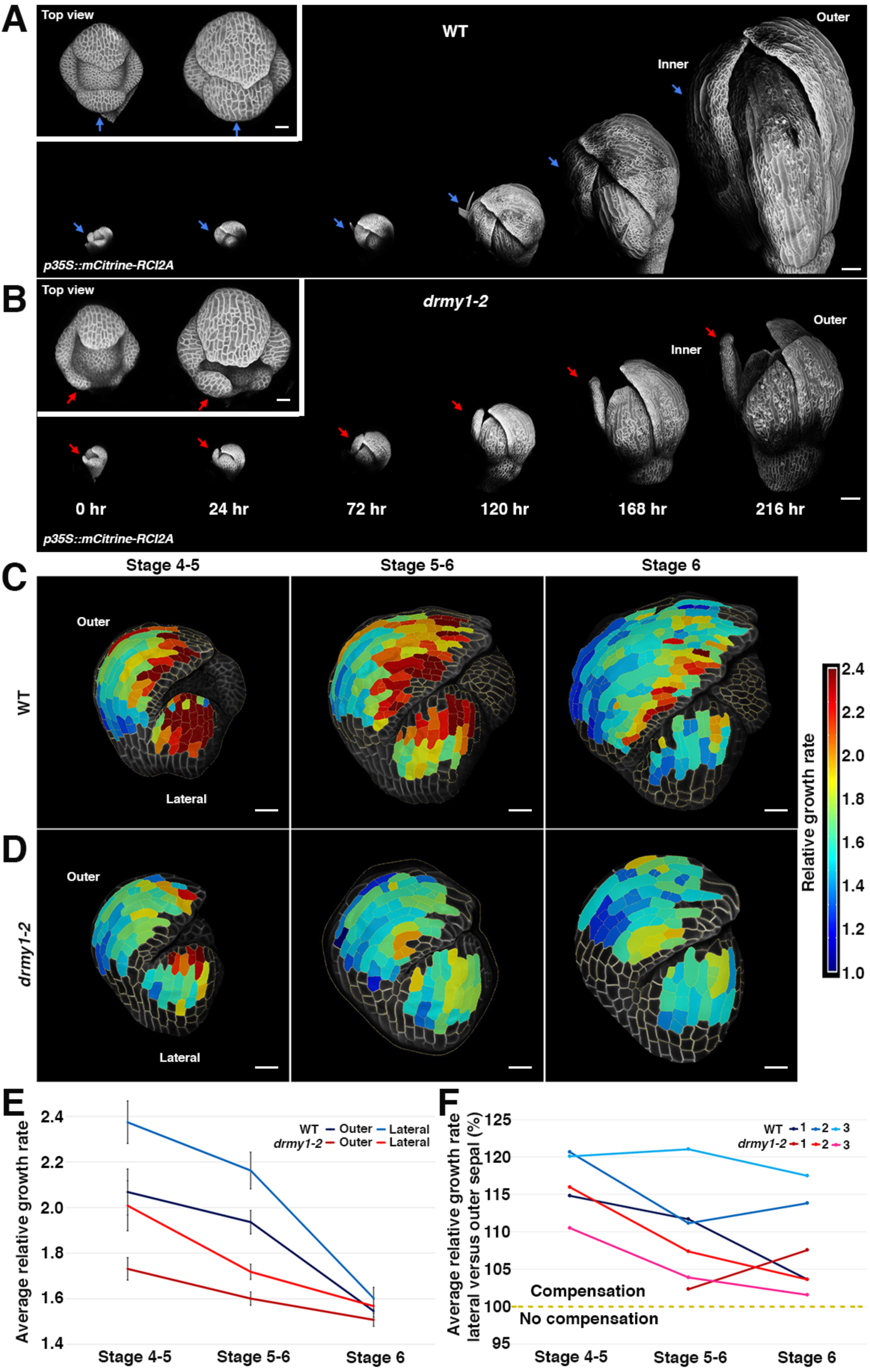
Sepals with delayed initiation in *drmy1-2* mutants remain smaller throughout development. (A and B) 24-hour live imaging of WT (A) and the *drmy1-2* mutant (B) flowers. The closed flower indicated the robust sepal size in WT while the *drmy1-2* flower remained open due to variable sepal size. n = 3; Inset: the top view of the flowers for the first two time points. Arrows: inner sepals; Scale bars: 100 µm; Scale bars in insets: 20 µm. (C and D) 12-hour early stage (from stage 4 to stage 6) cellular growth heatmap for both WT (C) and *drmy1-2* (D) outer (left) and lateral (center) sepals. The sepal cellular growth rate was quantified from live imaging of sepals immediately following initiation. For the heatmap, red indicates high relative growth rate while blue indicates low relative growth rate. Relative growth rate is defined as final cell size divided by initial cell size. Segmented cells are outlined in dashed yellow and superimposed on the meshed surface where the cell plasma membrane images were projected (greyscale). Scale bars: 20 µm. (E) Average cell growth rate curves of these early stage outer and lateral sepals for both WT and *drmy1-2*. Error bars: standard errors of the mean. (F) Relative cellular growth rate of the lateral sepal versus the outer sepal in the same flower. The average cellular growth rate of the lateral sepal was divided by that of the neighboring outer sepal and then percent was calculated. The yellow dashed line at 100% represents equal growth rates. Compensation happens above 100% as the lateral sepal grows faster to catch up.

### Growth patterns in *drmy1-2* mutants weaken their ability to compensate for early defects

Hypothetically, we would expect a late-initiating sepal to compensate for its smaller size by growing faster so that it can catch up with the other sepals in the same flower. We thus quantified compensation by live imaging outer and lateral sepals in the same flower (Fig. 3C-D). We tracked cells and their resultant daughters, enabling us to measure cell growth and division rates. In wild type, lateral sepals initiated 12 hours after outer sepals, and we observed that cells in lateral sepals generally grew faster than cells in the neighboring outer sepals (about 110-120% the average growth rate of the outer sepal cells; Fig. 3F). In *drmy1-2* mutants, lateral sepals initiate 12 to 30 hours after outer sepals and grew about 105-115% the average growth rate of their neighboring outer sepal cells, which would not be sufficient to fully compensate for later initiation (Fig. 3F). In both outer and lateral sepals, *drmy1-2* cellular growth was slower than wild type (Fig. 3C-E and Fig. S3A-C). We also noticed that cell division was reduced in the *drmy1-2* sepals (Fig. S3D). These results indicate that these later stage *drmy1* sepals also exhibit growth defects. Accordingly, the growth patterns of *drmy1* sepals weakens their ability to compensate for their delayed initiation, perpetuating sepal size variability throughout development.

Previously, we have shown that spatiotemporal averaging of variable cell growth results in sepal size and shape robustness and that this process is disrupted in *vos1* ^14^. Spatiotemporal averaging occurred normally in the *drmy1-2* mutant during early stage growth, indicating that the loss of robustness was due to distinct mechanisms (Fig. S4).

### Weak and diffuse auxin responses in *drmy1-2* mutants correlate with delayed sepal initiation

We next investigated how DRMY1 regulates the timing of sepal initiation. In the flower, auxin induces cell wall loosening, promoting cell expansion and allowing the primordium to emerge ^18^. Before the primordium initiates or bulges, the first sign of the incipient primordium is a localized region of auxin signaling created by the polarized transport of auxin ^19–22^. We examined the auxin response reporter DR5 (*pDR5rev::3XVENUS-N7)* ^19, 23^. In wild-type floral meristems, we found that the positions of incipient sepal primordia were marked by the expression of DR5 before primordium initiation occurs (Fig. 4A). Consistent with the variably delayed primordium initiation in *drmy1-2*, expression of the DR5 auxin response reporter was weaker and more diffuse in *drmy1-2* mutants (Fig. 4A and S5A, quantified in Fig. 4C). Weaker and more diffuse DR5 fitted with higher stiffness and slower growth in *drmy1-*2 mutants. Still, sepal primordia emerged from regions of auxin signaling in *drmy1-2* mutants. Positions where auxin signaling reaches high enough levels to initiate primordia is determined by the polar localization of the auxin efflux carrier PINFORMED1 (PIN1) ^19, 22^. PIN1 protein continued to polarize in *drmy1-2* inflorescence meristems and early flowers, so the more diffuse auxin response could not be easily explained by a loss of PIN1 polarity (Fig. S5C and D). Consistent with a decrease in auxin signaling, *drmy1-2* mutant plants exhibited a number of additional phenotypes associated with auxin signaling mutants: enhanced bushiness of the plant, shorter plant stature ^24^, smaller root meristem ^25^, shorter roots and fewer lateral roots ^26^ (Fig. S5E-G). Together these data suggest auxin signaling/response is reduced and more diffuse in *drmy1* mutants, correlating with delayed sepal primordium initiation.

**Fig. 4.**
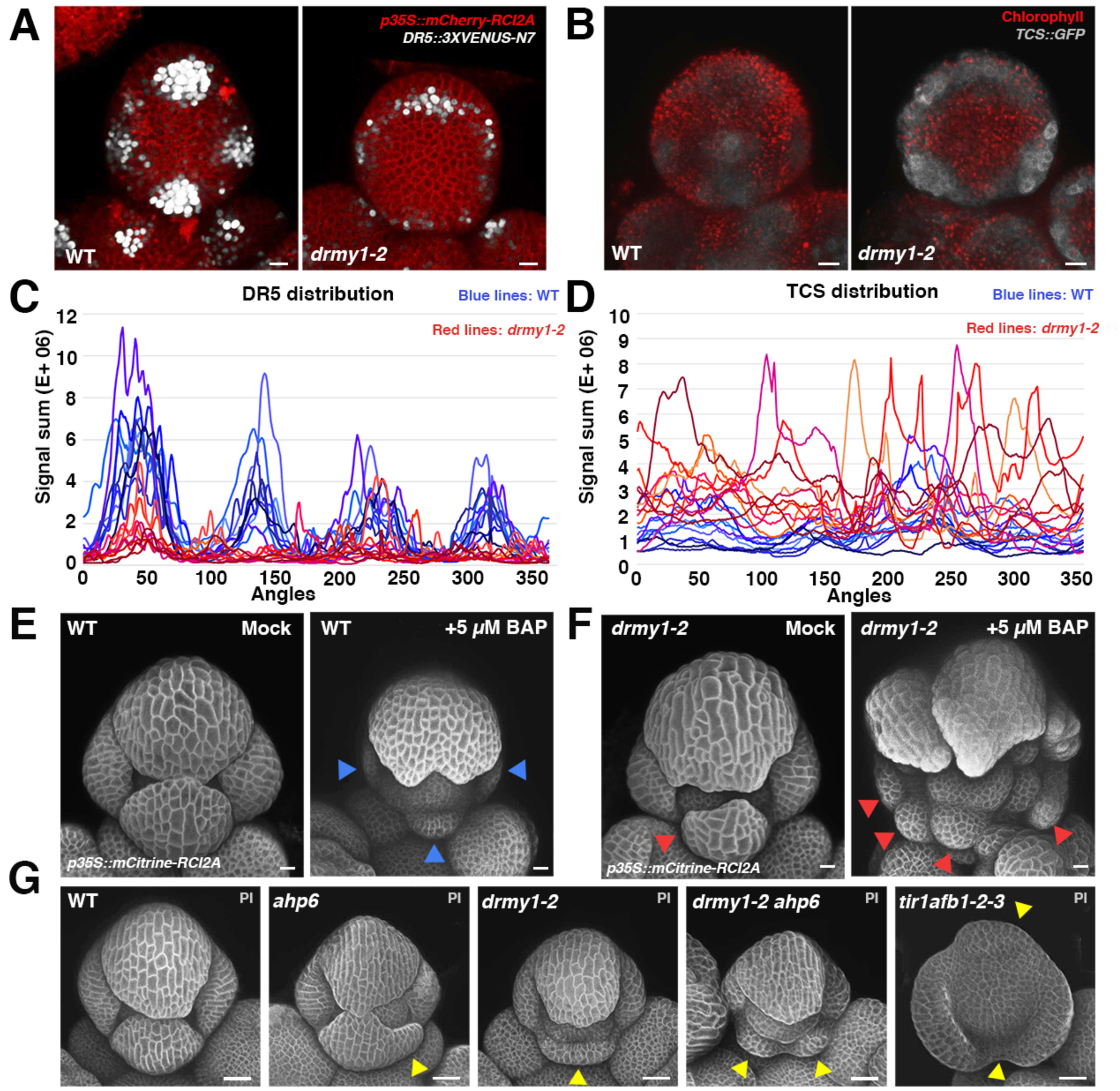
Restricted auxin and cytokinin signaling are required for robust sepal initiation. (A) Expression of the auxin response reporter DR5 (*DR5::3XVENUS-N7*, white) accumulates at the four incipient sepal initiation positions in WT. DR5 expression is lower and more diffuse in the *drmy1-2* mutant. *p35S::mCherry-RCI2A*: red, plasma membrane; Scale bars: 10 µm. (B) Expression of the cytokinin response reporter TCS (*pTCS::GFP*, green) accumulates at the four incipient sepal initiation positions in WT. TCS expression is enhanced and more diffuse in the *drmy1-2* mutant. Chlorophyll auto-fluorescence: red; Scale bars: 10 µm. (C) Quantification of DR5 signal in WT (blue curves) and *drmy1-2* (red curves) in stage 2 flowers when no sepals have initiated yet. Signal was quantified radially for the 360 degrees of the approximately circular flower meristem. The top-left region between the outer sepal and the lateral sepal was defined as angle 0. Angles increased in the counterclockwise direction and normalized signal values within bins of the size of 1° are plotted. n = 10 for both WT and *drmy1-2*. Note that four clear peaks of DR5 signal are present in WT. In *drmy1-2*, the outer sepal peak is evident, although weaker, and the remainder of the flower signal is relatively low without evident clusters. (D) Quantification of TCS signals in WT (blue curves) and *drmy1-2* (red curves) flowers at stage 2 when no sepals have initiated yet. n = 10 for both WT and *drmy1-2*. Note that four clusters of TCS signal are evident in WT, whereas in *drmy1-2*, TCS expression is higher and tends to surround the meristem. (E) Stage 6 flowers, where the sepals just close, from WT inflorescences cultured in 5 µM synthetic cytokinin BAP or mock media for 6 days. Blue arrowheads: delayed sepal initiation. Scale bars: 10 µm. (F) Stage 6 *drmy1-2* flowers from inflorescences cultured in 5µM BAP or mock media for 6 days. Red arrowheads: smaller sepals, indicating delayed sepal initiation. Scale bars: 10 µm. (G) The extent of disruption of auxin and cytokinin responses correlates with the degree of variability of sepal initiation timing. Mutation of *ahp6*, a cytokinin signaling inhibitor, has a very mild phenotype on its own, but enhances the *drmy1-2* sepal initiation phenotypes. Quadruple mutations of the auxin receptors *tir1 afb1-2-3* has a severely delayed sepal initiation phenotype. Cell walls stained with Propidium Iodide (PI) in grayscale. Yellow arrowheads: smaller sepal than normal, indicating the delayed sepal initiation. Scale bars: 50 µm.

### Strong and diffuse cytokinin responses in *drmy1-2* mutants correlate with delayed sepal initiation

Through its crosstalk with auxin, the plant hormone cytokinin controls the precise timing of flower primordium initiation within inflorescence meristems ^27, 28^. Therefore, we tested whether cytokinin signaling is involved in sepal primordium initiation and was altered in *drmy1-2* mutants using the cytokinin signaling reporter *pTCS::GFP* ^29^. In wild-type flowers, *pTCS::GFP* was expressed in the four incipient sepal positions, consistent with a role for cytokinin in primordium initiation. In the *drmy1-2* mutant, the expression of *pTCS::GFP* expanded and in some flowers formed a ring shape in the periphery of the floral meristem where the sepals should initiate (Fig. 4B and S5B, quantified in Fig. 4D). *drmy1-2* mutant plants also exhibited additional phenotypes associated with increased cytokinin signaling: disordered sequence and positions of flowers around the stem (phyllotaxy) ^27^, and enlarged inflorescence meristems ^30, 31^ (Fig. S5H-J).

### Auxin and cytokinin signaling patterns are required for robust timing of sepal initiation

Based on the hormone signaling reporters and hormone related phenotypes, cytokinin response increased and auxin response decreased in *drmy1-2* mutants. More importantly, the tight spatial localization of response reporters became more diffuse in *drmy1-2*. We therefore used three different ways to disrupt the auxin or cytokinin signaling and tested whether these disruptions could cause defects in the timing of sepal primordium initiation: increasing cytokinin, decreasing auxin signaling, and disrupting the crosstalk.

First, we tested whether broadly increasing cytokinin signaling was sufficient to delay sepal primordium initiation by externally applying cytokinin to whole floral meristems in wild type. We cultured dissected wild-type inflorescences in 5 µM cytokinin (BAP) media or mock media for 6 days. Cytokinin-treated flowers exhibited delayed and more variable sepal primordium initiation, which mimicked the phenotypes of *drmy1-2* mutants (Fig. 4E and Fig. S6A). We verified that this cytokinin treatment not only increased TCS signals but also made TCS expression more diffuse (Fig. S6E). As controls, the cytokinin receptor mutant *wol-1* was relatively insensitive to the treatment (Fig. S6F) and the TCS reporter remained unaffected when the inflorescence was treated with auxin (0.1, 1, and 20 µM NAA), confirming its specificity to cytokinin (Fig. S6G). Culturing *drmy1-2* mutant inflorescences in 5 µM cytokinin also enhanced the sepal initiation defects (Fig. 4F and Fig. S6B). This shows that the delay in organ initiation is not maximal in *drmy1*, and it further suggests that organ initiation delays are associated with disrupted cytokinin patterns.

Having shown that enhanced cytokinin signaling could delay sepal initiation, we then checked whether reducing auxin signaling throughout the flower was sufficient to delay sepal primordium initiation. Auxin signaling is inhibited in the auxin receptor quadruple mutant *tir1-1afb1-1afb2-1afb3-1* ^32^ and we observed similar, but more severe defects in the timing of sepal primordium initiation defects compared with *drmy1-2* (Fig. 4G). Thus, auxin signaling is also required for proper timing of sepal initiation.

Finally, we tested whether crosstalk between auxin and cytokinin is necessary for robust sepal initiation. We found that 5 µM cytokinin made the DR5 auxin response reporter’s signals more diffuse, as observed in *drmy1-2* mutants (Fig. S6C); this suggested that broader and increased cytokinin signaling may contribute to the diffuse DR5 auxin responses observed in *drmy1-2* mutants. However, we noted that DR5 expression levels did not decrease upon cytokinin treatment, in contrast with the *drmy1-2* mutant, implying that DRMY1 has a role in promoting auxin signaling. Furthermore, when treated with 5 µM cytokinin, the polarity pattern of PIN1-GFP appeared similar to that in *drmy1-2* mutants (Fig. S6D).

In the inflorescence meristem, high auxin at floral primordia positions activates MONOPTEROS (MP), which induces the expression of *Arabidopsis* HISTIDINE PHOSPHOTRANSFER PROTEIN 6 (AHP6). AHP6 acts as a cytokinin signaling inhibitor to define a brief period during which auxin and cytokinin signaling overlap to trigger primordium initiation ^27, 28^. In the *ahp6* mutant, initiation of sepal primordia was mildly affected compared to *drmy1-2* (Fig. 4G). Thus, DRMY1 plays a more prominent role to ensure the robustness of sepal primordium initiation than AHP6. In *ahp6 drmy1-2* double mutants, the delayed initiation phenotype was enhanced (Fig. 4G), suggesting that DRMY1 and AHP6 regulate robustness in primordium initiation synergistically. Taken together, our results suggest that DRMY1-dependent patterns of auxin and cytokinin signaling are critical for the robust temporal pattern of sepal initiation.

### Focused auxin and cytokinin signaling regions define zones of competence for sepal initiation

How do the spatial patterns of auxin and cytokinin signaling determine the temporal pattern of sepal initiation? To answer this question, we used live imaging to track expression of the DR5 auxin response reporter and the TCS cytokinin response reporter throughout the initiation of sepal primordia in developing flowers (Fig. 5A, Movie S3, S4 and S6). In wild-type incipient sepal primordia before initiation, the DR5 auxin response signal accumulated first in the outermost (L1) cell layer. Simultaneously, the TCS cytokinin response reporter was expressed directly below the DR5 signal. Both signals were restricted to incipient sepal positions. Over time, the DR5 signal invaded the inner cell layers (L2 and L3) leading to the coexistence of DR5 and TCS signals. After this overlap, we observed the outward bulges of primordium initiation. Then TCS and DR5 signals separated again. TCS signal remained at the base of the growing sepal, complementary to the DR5 signal which accumulated at the tip. In *drmy1-2* mutants, the invasion of DR5 signal into the inner cell layers was decreased and delayed at the inner sepal (Fig. 5B-C, Movie S5). Simultaneously, TCS expression was expanded around the periphery of the floral meristem and not limited to the incipient sepal positions in *drmy1-2* (Fig. 5D-E, Movie S7). Slowly the TCS signal resolved to a sharp domain of expression at the incipient sepal position in *drmy1-2* mutants (Fig. 5E, Movie S7). Once both the sharp TCS domain and the DR5 invasion were achieved, the *drmy1-2* inner sepal bulged out, indicating initiation. Although delayed, the focused domains of reporter expression were still eventually established in *drmy1-2* mutants. Our results suggest that establishing precisely localized and limited domains of both auxin and cytokinin signaling is required for sepal initiation, and that sepal initiation is variably delayed in *drmy1-2* until such precise domains can be established.

**Fig. 5.**
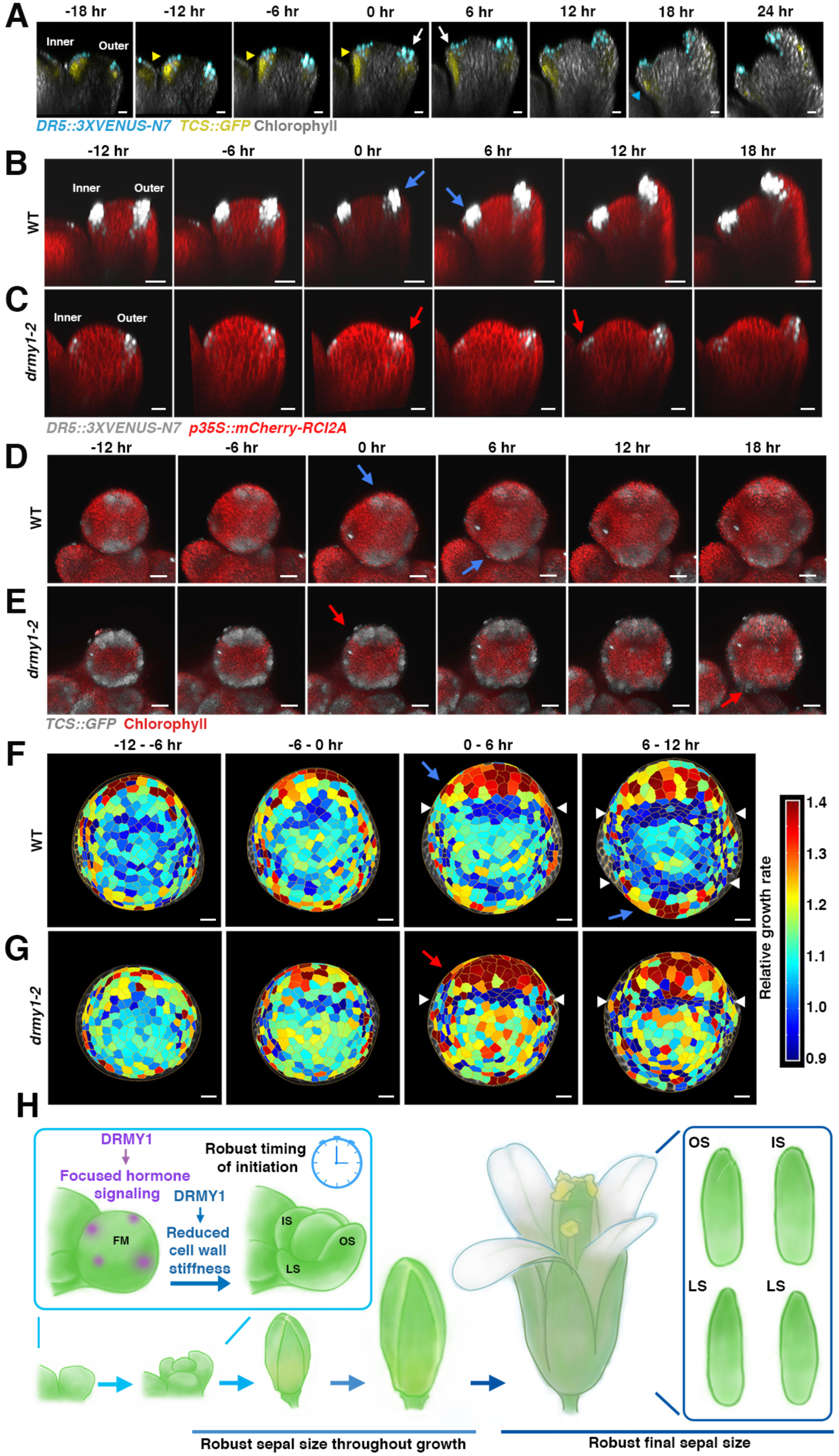
Spatiotemporal patterns of auxin and cytokinin signaling regulate the timing of sepal initiation. (A) Dual marker live imaging tracking DR5 (auxin, cyan nuclei) and TCS (cytokinin, yellow) signaling reporter expression throughout sepal initiation in WT. Longitudinal sections through the developing flower are shown with outer sepal on the right and inner sepal on the left. Chlorophyll (grey) outlines the morphology of the flowers. Yellow arrowheads: overlapping DR5 and TCS signals; Blue arrowheads: separation between DR5 and TCS; White arrow: sepal initiation event; Scale bar: 10 µm. (B and C) Live imaging of DR5 auxin signaling reporter (white) expression throughout sepal initiation in both WT (B) and the *drmy1-2* mutant (C). Longitudinal sections through developing flowers are shown with outer sepal on the right and inner sepal on the left. *p35S::mCherry-RCI2A*: red, plasma membrane; blue and red arrows indicate the sepal initiation. Note that the invasion of DR5 into inner layers is delayed of the inner *drmy1-2* sepal and is followed immediately by outgrowth. Scale bar: 20 µm. (D and E) Live imaging tracking TCS cytokinin signaling reporter expression (white) throughout sepal initiation in both WT (D) and the *drmy1-2* mutant (E). Top view of the developing flower shown with outer sepal on the top and inner sepal on the bottom. Blue and red arrows indicate sepal initiation. Chlorophyll autofluorescence: red; Scale bar: 20 µm.(F and G) Cellular growth heatmap throughout sepal initiation in WT (F) and *drmy1-2* (G). Top view of the flowers with the outer sepal at the top and the inner sepal at the bottom. For the heatmap, red indicates high relative growth rate while blue indicates low relative growth rate. Relative growth rate is defined as final cell size divided by initial cell size. White arrowheads: the band of cells with slower growth which specify the boundary between initiating sepals and the center of floral meristem. Blue and red arrows indicate the sepal initiation. Segmented cells are outlined in yellow. Scale bar: 10 µm. For all live imaging series, the time point when outer sepal primordia emerged were defined as time point 0. Images were staged relative to the timing of outer sepal initiation (0h) (H) DRMY1 ensures the focused hormone signaling and reduced cell wall stiffness during the sepal initiation process, thus making temporal sepal initiation pattern robust. The robust timing of sepal initiation is crucial for the sepal size robustness both throughout growth and at maturity.

### Tightly localized cell growth is associated with primordium initiation

Auxin and cytokinin regulate cellular growth ^18, 33, 34^. Since the spatiotemporal accumulation of hormone reporters was disrupted in *drmy1-2*, we also tested whether cell growth was affected during primordium emergence. The bulging of primordia requires both tightly localized regions of fast longitudinal growth at the periphery and slow latitudinal growth at the boundary between the primordium and the meristem center ^35, 36^. We analyzed the cellular growth rates and growth anisotropy of emerging primordia from live imaging, which were developmentally staged by the outer sepal morphology (Fig. S7E). In both wild type and *drmy1-2* mutants, before sepal initiation, cells grow heterogeneously without clear spatial pattern. Primordium initiation occurred with the appearance of a coordinated zone of fast-growing cells at the periphery and a tight band of slow growing cells at the boundary. For wild type, this switch to orchestrated spatial growth regions occurred at the inner sepal about 6 hours after the outer sepal (Fig. 5F, Fig. S7A), corresponding to the previous quantification of time intervals between sepal initiation (Fig. 2). For *drmy1-2*, although fast and slow growth regions began normally for the outer sepal, cellular growth within the inner sepal region remained heterogeneous 6 hours after the outer sepal initiation (Fig. 5G, Fig. S7B). As an independent test, we analyzed growth anisotropy. A switch from isotropic growth to highly anisotropic growth led to primordium initiation in both WT and mutant (Fig. S7C). In *drmy1-2*, the cellular growth at the regions where the inner sepal should initiate remained isotropic and randomly oriented during the time interval we analyzed (Fig. S7D). Thus, in *drmy1-2*, the inability to create tightly localized auxin and cytokinin signaling patterns coincided with stiffer cell walls and the inability to create tightly localized growth patterns, delaying initiation.

## Discussion

In this study, we report that DRMY1 ensures sepal size uniformity by coordinating the timing of sepal initiation (Fig. 5H). Because the *drmy1-2* mutant delays but does not block sepal initiation, it provides insights into the mechanisms controlling the timing of organ initiation. It is well established that the pattern of auxin accumulation sets the position of organ initiation ^19–22^. We observe that the sepal primordium does not emerge as soon as auxin signaling markers become apparent (Fig. 5A). Instead, stable focused regions of auxin and cytokinin signaling appear to define competency zones which give rise to tightly localized growth patterns required for organ initiation (Fig. 5H). In the case of auxin, a key event in the establishment of this focused region appears to be the invasion of auxin signaling into underlying cell layers, which later begins vascular development ^37^. Consistent with our results, this auxin invasion has been shown to stabilize the positions at which floral primordia form in inflorescence meristems ^37^. Auxin promotes growth through loosening the cell wall, and cell wall stiffness feeds back to regulate the polarity of the auxin efflux transporter PIN1^38, 39^. It remains for the future to determine how DRMY1 impinges on this feedback loop between cell wall stiffness, growth, and hormones.

Timing is important for developmental robustness in animals ^40^. For example, the robustness of somite size is generated by the precise timing of the somite segmentation clock ^41^. The duration of Bicoid exposure is also critical for patterning the Drosophila embryo ^42^. An implication of this work in *Arabidopsis* is that developmental timing of initiation can have cascading effects on organ size. In *drmy1-2* mutants, the late-initiating sepals remain smaller throughout development, so that sepal size remains variable. In sepals, uniformity of size is required throughout their growth to enclose the flower bud completely, creating a barrier with the external environment ^12^. Thus, traditional compensation mechanisms that delay maturation and termination of growth until the organs reach the right size, such as *dilp8* in *Drosophila* ^8, 9^, serve no purpose in sepals. Instead we see that wild-type lateral sepals, which emerge 12 hours after the outer sepals, compensate by growing faster to catch up. However, while wild-type lateral sepals close the flower bud, they remain slightly smaller even at maturity (Fig. S3F), suggesting that even in wild type, compensation may not be complete. In *drmy1-2*, the further delayed initiation of organ primordia, combined with a weaker ability to accelerate growth, perpetuates asymmetry in sepal sizes. We conclude that mechanisms ensuring precise timing of initiation make major contributions to robustness of organ size throughout development.

## Acknowledgments

We thank Fabrice Besnard, Anthony Bretscher, Joseph Cammarata, Kate Harline, Jessica McGory and Batthula Vijaya Lakshmi Vadde for comments on the manuscript. We thank Xiaoying Zhu for drawing the anatomical diagrams by hand. We thank Elliot Meyerowitz and Arnavaz Garda for sharing seeds for *DR5::3XVENUS-N7*/*PIN1::GFP* (Ler). We thank Teva Vernoux and Géraldine Brunoud for sharing seeds for *pTCS::GFP* (Col), *pPIN1::PIN1-GFP* (Col), and TCS DR5 (Col). We thank Fal Kateryna and Feng Zhao for teaching the protocol for PIN1 immunolocalization. We thank Fabrice Besnard for assisting with the phyllotaxy measurement.

## Author contributions

Conception and design for experiments: M.Z., A.H.K.R, V.M., A.B., O.H., R.S.S., and C.B.L. Isolation of *vos2* (*drmy1-2*) mutant: L.H. and A.H.K.R. Phenotypic analysis of *vos2* (*drmy1-2*) mutant: M.Z. and Z.W. Variability of organ shape analysis: C.B.L. SEM: M.Z. and A.H.K.R. Live imaging and analysis: M.Z. and W.C. AFM: M.Z and S.B. MorphoGraphX plug-in developed for DR5 and TCS quantification: S.S. and R.S.S. Computational analysis of spatial and temporal variability of growth: S.T. and C.B.L. RNA-seq analysis: E.M.S., M.Z., and A.H.K.R. Writing of the manuscript: M.Z. and A.H.K.R. Revising and editing of the manuscript: M.Z., W.C., V.M., L.H., S.B., S.S., E.M.S., S.T., R.S.S, C.B.L., O.H., A.B., and A.H.K.R.

## Data and materials availability

RNA-seq data are available at NCBI BioProject PRJNA564625. All other data is available in the main text or the supplementary materials. Materials are available by request.

## Supplementary Materials

### Materials and Methods

#### Plant Growth Conditions

Most of the plants were grown in soil under 24-hr fluorescent light conditions (∼100 µmol m^−2^ s^−1^) at 22°C in Percival growth chambers. Plants used for live imaging were grown in soil under 16h light (∼100 µmol m^−2^ s−^1^) / 8h dark long day conditions at 22°C in a Percival growth chamber to produce relatively larger shoot apical meristems (SAMs). These different growth conditions did not affect the *drmy1* mutant phenotypes we reported here. All seeds were sown on Lambert Mix LM-111 soil and cold-stratified at 4°C for 2 days. Plants used for root phenotyping were grown in ½ Murashige and Skoog (MS) media (pH5.7, 1% agar) under 24-hr fluorescent light conditions (∼100 µmol m^−2^ s^−1^) at 22°C in Percival growth chambers.

#### Mutations and genotyping

In this study, we primarily use *Arabidopsis thaliana* accession Col-0 as the wild-type plants. As described in Hong et al., 2016 ^14^, *<u>v</u>ariable <u>o</u>rgan <u>s</u>ize and shape* (*vos)* mutants were isolated from an M2 population of ethyl methanesulfonate (EMS) mutagenized Col-0. The *vos2* (*drmy1-2*) mutant was back crossed to Col-0 three times to segregate unrelated mutations before further characterization. The *vos2* (*drmy1-2*) mutant was then crossed with an *Arabidopsis* Landsberg *erecta* accession plant to generate a mapping population. The *vos2* (*drmy1-2*) mutated gene was identified through map-based cloning following the standard procedure described in ^43^. The *vos2* (*drmy1-2*) mutation was mapped to an interval containing 97 genes on chromosome 1 between 21.3M and 21.7M. The *vos2* (*drmy1-2*) mutant contains a G to A mutation at the junction between the third intron and the fourth exon within the *<u>D</u>EVELOPMENT <u>R</u>ELATED <u>M</u>YB-LIKE 1 (DRMY1,* AT1G58220) gene, which was predicted to cause splicing defects that were later verified experimentally. The *drmy1-2* G to A point mutation was genotyped through PCR amplification with the dCAPs (Neff et al., 2002) primers oMZ113 and oMZ114 (sequences in Table S1), followed by the digestion of PCR products with DdeI to produce 74-bp WT products and 100-bp mutant products. We crossed *vos2* (*drmy1-2*) with *drmy1-1* (with a T-DNA insertion in the first intron; SALK012746, ^15^) to test for allelism. The resulting F1 exhibited the *drmy1* mutant phenotypes indicating these mutations failed to complement and are allelic. To verify the splicing defects, the mRNA was extracted from the *drmy1-2* mutant inflorescences, followed by RT-PCR to generate cDNAs. Mutated *DRMY1* CDS was amplified with primers listed in Table S1 and then inserted into *pENTR-D-TOPO* vectors (Invitrogen). The resultant plasmids were then purified and sequenced with commercial primer M13F.

#### Flower staging

All the flower staging was based on ^44^.

#### Photographing of flowers, inflorescences, and whole plants

Stage 12 and 15 flowers were dissected from the primary branches and put onto black papers. Primary inflorescences were dissected, and their stems were captured with tweezers for positioning. Black paper was also put under the tweezers. A Canon Powershot A640 camera on a Zeiss Stemi 2000-C stereomicroscope was then used to photograph the flowers and inflorescences with black background. Whole plants of both WT and *drmy1-2* were taken from the soil and washed. They were then put on the black cloth. Photographs of whole plants were then taken with Canon Powershot SD1300 camera.

#### Sepal area and shape analysis

Full-size, mature, stage 15 sepals were dissected from the flowers for analysis. For sepal area and shape variability comparison between wild-type and *drmy1-2* mutants, we selected the 10th to 25th flowers from the main branch (primary inflorescence) because they are relatively consistent in wild type as confirmed in ^14^. For quantification of the mean and variance of the four sepals within a single flower, we dissected sepals throughout development from the earliest flower we could start (ranging from the second one to the fifth one) to the latest flower we could get (ranging from the 50th one to the 65th one) on the main branch. The sepal contour extraction analysis was done as described in ^14^. Briefly, the dissected sepals were flattened between slides and photographed with Canon Powershot A640 camera on a Zeiss Stemi 2000-C stereomicroscope. Custom python scripts were then used to extract the contour from each sepal photo and quantify the area and shape variability (scripts available in the supplementary material Data S1 of ^14^).The coefficient of variation (CV) of sepal areas was calculated by dividing the standard deviation by the mean of four sepal areas within a flower.

#### Meristem size analysis

The confocal image z-stacks of WT and *drmy1-2* inflorescences were collected with the inflorescence meristem well exposed through dissection. The LSM images were then converted to TIFF format with FIJI. The TIFF files were loaded into MorphoGraphX and oriented such that the center of the inflorescence meristem faced straight up. Screenshots were saved and loaded into FIJI again. For each meristem, the biggest circle was drawn to include the inflorescence meristem without capturing any emerged flowers. The areas of the circle were then measure and converted to square microns.

#### Phyllotaxy analysis

The phyllotaxy measurement and analysis was done followed the procedure described in Peaucelle et al., 2007 ^45^ and Besnard et al., 2014 ^27^. Briefly, WT and *drmy1-2* plants were grown in parallel. When the plants had more than 40 stage 17 siliques, the plants were attached to the device as shown in Peaucelle et al., 2007 Figure S1A. Divergence angles were then measured and recorded into the excel file. R functions (provided by Fabrice Besnard) were then used for analysis and generating plots.

#### Permutation test to confirm the difference between two populations

We used the permutation test to determine whether size distributions of sepals were significantly different between WT and *drmy1-2*. The permutation test does not require the knowledge of the underlying distribution functions. The permutation test was done as described in ^14^.

#### Scanning electron Microscopy (SEM) observation

SEM was performed with a Leica 440 through the Cornell center for Materials Research Shared Facilities (Supported through NSF MRSEC program DMR-1719875) as described in ^46^.

#### Live imaging of sepal initiation and growth

½ MS media containing 1% sucrose, 0.25x vitamin mix, 1µL/mL plant preservative mixture and 1% agarose (Recipe modified from ^47^) was poured into the small Petri dishes (Fisher 60mm • 15mm) for positioning inflorescences and supporting growth. In this paper, we used two different methods to dissect and position the *Arabidopsis* inflorescences for live imaging. The first method, viewing the inflorescence from the side, was modified from ^46^. First, we removed the MS media from half of the Petri dish to create space for the inflorescence. Inflorescences of plants expressing the plasma membrane markers *pLH13* (*35S::mCitrine-RCI2A*, yellow plasma membrane marker;^48^) were dissected with a Dumont tweezer (Electron Microscopy Sciences, style5, Cat #72701-D).

Overlying older flowers from one side were removed to expose the stage 4 flowers. The inflorescences were taped to a cover slip to ensure the correct orientation. The cover slip was then positioned in the Petri dish with the base of inflorescence stem inserted into the MS media. We then taped the Petri dish to the sides of the Percival growth chambers with inflorescence vertical and the bottom of the dish facing outwards to avoid growth bending. This method was mainly used for imaging the early stage lateral and outer sepal development.

In the second method, we imaged the inflorescence from the top, similar to the method reported in^47^. Primary inflorescences containing *p35S::mCitrine-RCI2A* (pLH13), or *DR5:: 3XVENUS-N7* (Auxin response reporter ^19^), or *pTCS::GFP* (Cytokinin response reporter ^49^), or both *DR5::3XVENUS-N7* and *p35S::mCherry-RCI2A* (pMZ11, LR reaction between pENTR with mCherry-RCI2A (pHM52) and pB7WG2 (destination vector with p35S), red plasma membrane marker), or both *DR5::3XVENUS-N7* and *pTCS::GFP* were dissected with tweezers and then the stem was inserted into the MS media positioning the inflorescence upright. Further dissection with the tip of the tweezer or a needle was done to remove all the flowers older than stage 4. This method was used for imaging the initiation of sepal primordia and the late stage abaxial sepal development.

After dissection and positioning, the petri dishes were kept in the growth chamber at least 6 hr for plant recovery before live imaging. The chosen flowers were imaged every 6 hr (sepal initiation), 12 hr or 24 hr (organ growth) with a Zeiss 710 confocal microscope. For long-term live imaging which lasted for more than 6 days, we transferred the inflorescences from the old petri dishes to newly made fresh ones for keeping active growth every three days. Before imaging, the inflorescences were immersed in the water for at least 10 minutes and 20x Plan-Apochromat NA 1.0 water immersion objective was used for imaging. The wave lengths for the excitation and emission for the florescent proteins were indicated in Table S2. The depth of z-sections was set to 0.5 µm (live imaging for sepal growth) or 1 µm (live imaging for sepal initiation or reporter patterning). The resultant LSM files were converted and cropped with FIJI. MorphoGraphX was used for visualization of the spatial distribution of florescent signals and creation of digital longitudinal sections.

#### Image processing for growth quantification

Imaging processing and growth quantification were performed as described in ^14^. Briefly, the confocal stacks collected from the live imaging of sepal growth were converted from LSM format to TIFF format with FIJI. The images were then imported into MorphoGraphX ^50^. Sample surfaces were detected and meshes of the surface were generated. Fluorescent signals were then projected onto the meshes and cells were segmented on the meshes. For cellular growth, cell lineages were defined manually and cell area between different time points were compared to quantify growth rates. Heatmaps were generated to visualize the areal growth rate and the values for the growth rate of each cell were then exported to CSV files. Graphs of growth trends were generated with the analysis of the CSV files in Microsoft Excel. The cell division heatmap was also generated based on the cell lineages, presenting how many daughter cells originated from one mother cell. The analysis of Principal Directions of Growth (PDG) was also done with MorphoGraphX, following the user manual. Briefly, the two meshes of two different time points were loaded together with the parent label. Check correspondence was done to make sure there were no errors of cell junctions. The growth directions were then computed and “Aniso (StrainMax versus StrainMin)” and “StrainMax” were visualized on the second time point mesh.

#### Analysis of spatiotemporal variability in the growth of cell area

The analysis of spatiotemporal variability in the growth of cell area was done as described in ^14^. Briefly, the cellular growth rates were quantified with MorphoGraphX. To calculate the local spatial variability, the areal growth rates for the cell of interest and its surrounding cells were defined and calculated with the V_area_ equation. To calculate the temporal variation, the areal growth rates for the cell of interest at one time frame and the following time frame were defined and calculated with the D_area_ equation. Both V_area_ and D_area_ were calculated for WT and *drmy1-2* flowers at the same stages and plotted.

#### Gaussian curvature measurement

The LSM flies from live imaging were first converted to TIFF files with FIJI. The TIFF files were then loaded in MorphoGraphX on Stack1. The following procedures performed: Gaussian blur stack 3 times (X/Y/Z sigma (µm) = 1); Edge detection (Threshold = 7000, Multiplier = 2.0, Adapt Factor = 0.3, Fill Value = 30000); Marching cube surface (Cube size (um) = 8, Threshold = 20000); Subdivide mesh 2 times; Smooth mesh (Passes = 5) 2 or 3 times; Project mesh curvature (Output = blank, Type = Gaussian, Neighborhood (um) = 10, Autoscale = no, Min Curv = −0.002, Max Curv = 0.002, Autoscale percentile = 85); Save the screenshot to JPEG images.

#### Imaging of reporter lines

Primary branches of the reporter line plants (*DR5::3XVENUS-N7*, *pTCS::GFP*, *PIN1-GFP*, *pDRMY1::3XVENUS-N7*, *pDRMY1::DRMY1-mCitrine*, *p35S::mCitrine-RCI2A*) were dissected with tweezers and inserted upright into ½ MS media (containing 1% sucrose, 0. 25x vitamin mix and 1% Agar) poured into small petri dishes. The samples were immersed in the water for 30min and then further dissected with the tip of tweezers to remove all unnecessary flowers. After dissection, the inflorescences were put in the growth chamber for 6 hours for recovery and then imaged with 20x Plan-Apochromat NA 1.0 water immersion objective on a Zeiss 710 confocal microscope.

The seedlings of both WT and *drmy1-2* with *p35S::mCitrine-RCI2A* were grown in ½ MS media (containing 1% sucrose, 0.25x vitamin mix and 1% agar) for around 5 days with Petri dishes placed vertically. The seedlings were then well positioned into waterdrops loaded in advance on the slides. After putting on the cover slip, the roots were imaged with 20x Plan-Apochromat NA 1.0 water immersion objective on the Zeiss 710 confocal microscope. The excitation and emission wavelength for the fluorescent proteins are indicated in Table S2.

#### Cytokinin (BAP) and auxin (NAA) treatments

Primary inflorescences containing target reporters were dissected and inserted into ½ MS media coated small petri dishes. The inflorescences were then put into the chamber for at least 6 hours for recovery. Cytokinin or auxin containing ½ MS media were made following the ½ MS media recipe mentioned before, with specific volumes of the 0.5g/L synthetic cytokinin (BAP) stock solution (kept in −20°C) or 5mM/L synthetic auxin (NAA) stock solution (kept in 4°C) for specific concentrations. The inflorescences were then transferred to the cytokinin or auxin containing ½ MS media for the external cytokinin or auxin treatments. 20x Plan-Apochromat NA 1.0 water immersion objective on the Zeiss 710 confocal microscope was again used for image collection. For the BAP treatment, *pTCS::GFP* reporter lines were used as the positive control. The cytokinin receptor mutant *wol* was used as the negative control as they were relatively insensitive to the BAP treatment.

#### PIN1 immunolocalization

PIN1 immunolocalization was done with the modified procedure described in ^51^. Briefly, the primary branches of WT and *drmy1-2* plants were dissected with tweezers and inserted upright into ½ MS media (containing 1% sucrose, 0.25 x vitamin mix and 1% agar) poured into small petri dishes. The meristem was fixed in FAA (3.7% formaldehyde, 5% acetic acid, 50% ethanol) for 1 hour. Ethanol gradient treatment was used to remove chlorophyll. Cocktail enzyme treatment helped to puncture the cell wall to allow absorption of antibodies. After blocking with 3% BSA, PIN1 antibody (PIN AP20, Santa Cruz Biotechnology) was applied to the samples, followed by the treatment of the secondary antibody (anti goat 488). The inflorescences were then mounted upright into 1% agarose medium and imaged with a confocal microscope.

#### Atomic force microscopy (AFM) and data analysis

To prevent vibrations, dissected meristems were inserted vertically into a ½ MS media (containing 1% sucrose, 0.25 x vitamin mix, 1% agar, pH 5.7) coating a 60 mm Petri dish (Falcon 60 mm • 15 mm, Corning Ref. 351007). In order to better stabilize the sample against movements/vibrations that can occur during indentation measurements, 0.8% low melting agarose melted in the water was added in order to make a dome around the sample, covering all but the top of the inflorescence meristem.

AFM experiments were performed on a stand-alone JPK Nanowizard III microscope, driven by JPK Nanowizard software 6.0. The acquisitions were done in Quantitative Imaging mode (QI). The experiments have been performed in Milli-Q purified (MQ) water, that was added into the petri dish 30 min to 1 hour before the beginning of the measurement to maintain meristem hydration. We used a silica spherical tip (Special Development SD-sphere-NCH, Nanosensors) mounted on a silicon cantilever with a nominal force constant of 42 N/m, and a radius of 400 nm. Scan size was generally of 50 µm with pixel size of 500 nm (e.g. 100 pixels for 50 µm scan size).

The applied trigger force was of 1 µN, a force corresponding to 200-300 nm of maximum indentation, in order to indent the cell wall only ^52, 53^. The ramp size was of 2 µm (1000 data points per curve), approach speed of 100 µm/s, and retract speed of 100 µm/s.

Cantilever calibration was performed following the standard thermal noise method. We measured the deflection sensitivity by doing a linear fit of the contact part of a force curve acquired on a sapphire sample in MQ water. Then, we determined the spring constant by acquiring the thermal noise spectrum of the cantilever and fitting the first normal mode peak using a single harmonic oscillator model. The same tip was used for several experiments in different days, as long as possible. In order to reduce the offsets in force that can be introduced by each new calibration, especially by the measurement of the deflection sensitivity, we followed the SNAP protocol developed by Schillers et al. ^54^. Since our cantilevers have not been independently calibrated by the producer, we consider the spring constant value found at the first calibration as the reference value. In the following calibrations, we correct the deflection sensitivity following SNAP, in order to obtain the same spring constant each time by thermal tune analysis. Then, the new deflection sensitivity and the reference spring constant will be set in the instrument’s software.

Data analysis was done using JPK Data Processing software 6.0. Force vs height curves were first flattened by removing the result of a linear fit done over a portion of the non-contact part (baseline), in order to set this part to 0 force. A first estimation of the point of contact (POC), was obtained considering the first point crossing the 0 of forces, starting from the end of the approach curve (i.e. trigger force position). The force vs. tip-sample distance was then obtained calculating a new axis of distances as Height [m] – tip deflection d [m]. Young’s modulus was obtained by fitting the entire force vs tip-sample distance curve with a Hertz model (as it is called in the JPK Data Processing software). The equation used for fitting has been derived as described in Sneddon, 1965^55^:

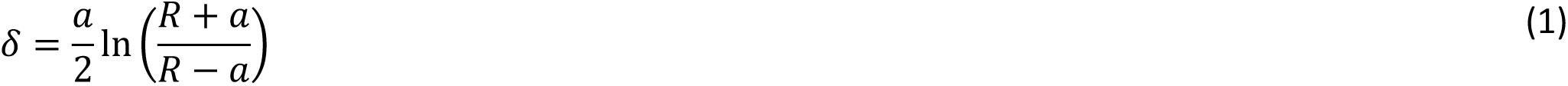

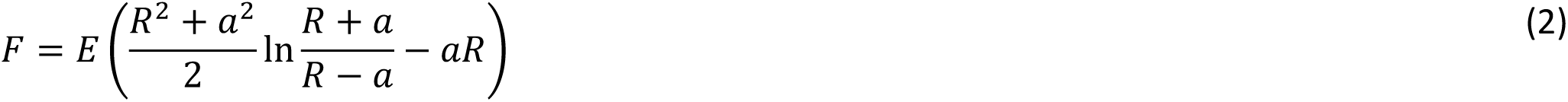

where R is the tip radius, δ and F the obtained indentation and the applied load, the radius of the contact area and E the reduced modulus = E* / (1 – ν^2^), ν being Poisson’s ratio, E being referred to as simply Young’s modulus in this study. (Note the typo in Eq. (6.15) of Sneddon, 1965 for F – there is an extra factor 1/2 in front of the −aR term.)

In order to express F as a function of δ, we can expand it in powers of δ/R finding:

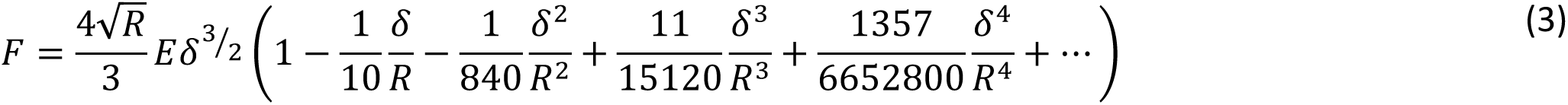

Generally, in AFM measurements, the Hertz model is only the first term of this expansion, which is named ‘Paraboloid’ in JPK Data Processing. The difference between the results provided by the two models, becomes important only when δ ≈ R, so in any case at the limit of the applicability of both formulae.

For our analysis, we used a tip radius R of 400 nm and a Poisson’s ratio ν of 0.5 (as it is conventionally set for biological materials), where the Young’s modulus, the POC and an offset in force were kept as free parameters of the fit. The same analysis protocol was used on approach and retract curves.

On samples like meristems, the contact between the indenting tip and the surface is generally not normal. As discussed by Routier-Kierzkowska ^56^ this will lead to artifacts in the determination of Young’s modulus. In order to remove or reduce the effect of the local slope on our calculations, we adapted a formula from the previously cited paper and calculated the normal Young’s modulus as:

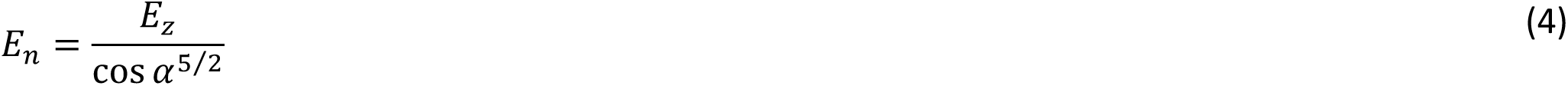

where α is the angle between the normal to the surface and the z direction and n is the normal to the surface.

#### Osmotic treatments measuring sepal stiffness

The osmotic treatments were done as described in ^14^. Briefly, the inflorescences containing plasma membrane markers *p35S::mCitrine-RCI2A* (*pLH13*) were dissected from the main branch with one stage 8 flower exposed and inserted upright into small Petri dishes containing ½ MS media. The inflorescences were then incubated in water for osmotic pressure saturation. 0.1% PI was used to visualize the cell walls. The inflorescences were imaged on the confocal microscope before the osmotic treatment (with wave lengths as indicated in Table2) The water was then removed and a 0.4M NaCl solution was used to plasmolyze the cells. The separation between the cell wall (stained with PI) and the cell membrane (labeled with mCitrine) indicated that plasmolysis was successful. The confocal images were taken after 30 min salt treatments with the sample still immersed in the salt solution. Cell areas were quantified before and after salt treatment in MorphoGraphX and the change (shrinkage) in cell area indicated the (elastic) stiffness of the cell walls (little or no change occurs in stiff walls while softer walls shrink).

#### Transgenic plants

The *DRMY1* gene promoter with 5’ UTR (1,535 bp before the start codon) and the terminator with 3’ UTR (331 bp after the stop codon) were PCR amplified with primers listed in the Table S1. These two pieces were fused with overlap PCR with PFU DNA polymerase. The PCR product was A tailed with Taq DNA polymerase. With the help of this overhanging A, the overlap PCR product was then ligated into *pGEM-T easy* (Promega) to generate *pMZ2*. The Gateway conversion cassette was PCR amplified and digested with restriction enzymes AscI and KpnI and then ligated into *pMZ2* between the *DRMY1* promoter and terminator to make *pMZ3*. The resulting *pMZ3* plasmid and binary vector *pMOA36* were digested with NotI and ligated together to make *pMZ4*. The final *pMZ4* Gateway destination vector includes the *DRMY1* promoter, a gateway cassette, and the *DRMY1* terminator. The *DRMY1* gene CDS was PCR amplified with primers listed in the Table S1. The *DRMY-mCitrine* fusion was created by amplifying the CDS of *DRMY1* and the CDS of *mCitrine* with primers listed in Table S1. The *DRMY1-mCitrine* fusion was created through overlap PCR with a nine-alanine linker in the middle. Each of these PCR products was purified and cloned into *pENTR/D-TOPO* vectors (Invitrogen). The resultant vectors were LR combined into the destination vector *pMZ4* to generate the final constructs used in this paper: *pDRMY1::DRMY1* (*pMZ6*) and *pDRMY1::DRMY1-mCitrine* (*pMZ18*)). All of the final constructs were verified by sequencing and transformed into the *drmy1-2* mutants by Agrobacterium-mediated floral dipping. All the T1 plants were grown in soil for about 10 days and then selected by spraying with 100 µg/mL Basta. The surviving plants were then checked for sepal phenotypes.

#### Generating and analyzing RNA-seq data

For sample collection, the inflorescences were dissected under a stereomicroscope and collected into Eppendorf tubes with liquid nitrogen. Both WT and *drmy1-2* inflorescences were dissected to remove flowers older than stage 8. Three biological replicates were collected, each replicate combining different individual plants. RNA extraction was done using a commercial kit (RNeasy Plant Mini Kit, Qiagen, CAT NO. 74904) following the manual.

The cDNAs used for qPCR were generated by reverse transcription from inflorescence RNA samples (DNase treatment, first strand synthesis and second strand synthesis). The qPCR was performed following the manual of the Roche LightCycler® 480 system. Briefly, synthesized cDNA, primers and water were mixed with Roche CyberGreen Master Mix in the Temp Plate 384-well full skirt PCR plates (USA Scientific). The PCR plate was then loaded into Roche 480 machine and the experimental data was analyzed with the supporting software.

RNA-seq library preparation was performed as in ^57^ with modifications. Briefly, mRNA was isolated with Dynabeads oligo dT beads (Invitrogen Cat# 61006) and fragmented in first strand buffer at 94° C for 6 minutes. First strand cDNA was synthesized with random hexamers (Invitrogen Cat# 48190011) using Superscript II (Invitrogen Cat#18064014) and the second strand was synthesized with DNA Pol I (Fermentas Cat #EP0041) and RNAse H (Invitrogen Cat# 18021071). End repair was conducted with NEBNext End repair enzyme Mix (NEB Cat# E6050S) and Klenow DNA polymerase (NEB Cat# M0210S), A tailing with Klenow 3’ to 5’ exonuclease (Fermentas Cat# EP0421), and ligation of NEB NEXT adapters (NEB Cat# E7335L) with Mighty Mix Ligase (Clonetech TAK6023). The library was size-selected using Agencourt AmPure Beads (Beckman Coulter A63880). PCR enrichment and barcoding were conducted with NEBNext Multiplex Oligos for Illumina index primers (NEB Cat# E7335L) for 15 cycles using Phusion polymerase (NEB Cat# M0530L). The entire library was run on a 1.2% agarose gel and size-selected (about 200– 500 bp) to remove adapter dimers. RNA sequencing was done with the service HISEQ 2500 rapid run (run length 50 RR, single barcode) or NextSeq 500 (run length 75, single barcode) provided by Cornell Genomic facility. Reads are available at NCBI BioProject PRJNA564625.

We analyzed our RNA-seq reads and annotated genes somewhat as in ^58^, but with a number of updated procedures described here. For quality-filtering of RNA-seq reads, we first used the Perl script *quality_trim_fastq.pl* (available at https://github.com/SchwarzEM/ems_perl/blob/master/ *illumina/quality_trim_fastq.pl*) to remove reads that failed CHASTITY and trim the last nucleotide off our raw reads; *quality_trim_fastq.pl* was run with the arguments ‘-*q 33 −u 50*’ (for replicates 1 and 2 of each genotype) or ‘-*q 33 -u 84*’ (for replicate 3 of each genotype). We then used Trimmomatic 0.36 ^59^ to remove adapter sequences and remove unreliable reads. Trimmomatic was run with the arguments *’SE-phred33 […] ILLUMINACLIP:[…]/sepal_adapters_09dec2016. trimmomatic.fa:2:30:10 LEADING:3 TRAILING:3 SLIDINGWINDOW:4:15 MINLEN:[50 or 84]*’. Adaptor sequences (*sepal_adapters_09dec2016.trimmomatic.fa*) are given in Table S3. After quality trimming, replicates had 6.5 to 25.6 million reads (Data S1). These trimmed and quality-filtered reads were permanently archived in the NCBI Sequence Read Archive (SRA) with their different read lengths between batches (50 nt for replicates 1 and 2 in an earlier batch; 84 nt for replicate 3 in a later batch). However, before mapping reads to transcripts for gene expression analysis, we further trimmed the 84-nt reads to 50 nt lengths with *quality_trim_fastq.pl* (arguments, ‘-*q 33 -u 50*’). By enforcing equal read lengths, we expected to lessen artifactual differences in observed gene expression that might otherwise arise from unequally long RNA-seq reads ^60^.

We mapped RNA-seq reads to cDNA sequences of *A. thaliana* genes (i.e., coding sequences [CDSes] plus flanking 5’ and 3’ UTR sequences) that were downloaded from http://www.arabidopsis.org/download_files/Sequences/TAIR10_blastsets/TAIR10_cdna_20101214_updated, along with a GFP coding sequence previously used in ^58^. We used Salmon 0.14.1 ^61^ to index the cDNA sequences and determine levels of gene expression for each RNA-seq replicate in transcripts per million (TPM). For indexing cDNA, we used the Salmon arguments *’--no-version-check index -k 31 --perfectHash --type quasi --keepDuplicates*’; for computing gene expression, we used the arguments ‘*--no-version-check quant --seqBias --gcBias --validateMappings --libType A --geneMap TAIR10_cdna_20101214_updated_w_ GFP.tx2gene.tsv.txt --numBootstraps 200*’. All replicates had mapping rates of 92.6% to 95.3% (Data S1).

To compare changes of gene expression between genotypes and identify which changes were statistically significant, we used DESeq2 version 1.18.1 ^62^, downloaded and run as a Bioconda package in its own environment (https://bioconda.github.io/recipes/bioconductor-deseq2/ *README.html*), with batch corrections for replicate 3 of each genotype (which were grown at a different time from replicates 1 and 2 of each genotype).

Gene annotations were produced as follows. Gene names, aliases, and functional descriptions, were downloaded from TAIR on 9/15/2019 via TAIR’s bulk download Web portal (*https://* www.arabidopsis.org/tools/bulk/genes/index.jsp) and tabulated with *tabulate_tair_names_ 15sep2019.pl* (https://github.com/SchwarzEM/ems_perl/blob/master/arabidopsis/tabulate_tair_names_15sep2019.pl). Protein sizes (full range or maximum) were extracted from the TAIR10 proteome (downloaded from http://www.arabidopsis.org/download_files/Sequences/TAIR10_blastsets/TAIR10_pep_20101214_updated; version dated 4/16/2012) with *tabulate_tairprot_sizes.pl* (https://github.com/SchwarzEM/ems_perl/blob/master/arabidopsis/tabulate_tairprot_sizes.pl). Signal and transmembrane sequences were predicted with Phobius 1.01 ^63^ and tabulated with *tabulate_phobius_hits.pl* (https://github.com/SchwarzEM/ems_perl/blob/master/Hco_Acey/tabulate_phobius_hits.pl). Coiled-coiled domains were predicted with ncoils ^64^ and tabulated with *tabulate_ncoils_x.fa.pl* (https://github.com/SchwarzEM/ems_perl/blob/master/ *Hco_Acey/tabulate_ncoils_x.fa.pl*). Low-complexity protein regions were predicted with pseg ^65^ and tabulated with *summarize_psegs_16jan2018.pl* (https://github.com/SchwarzEM/ems_perl/ *blob/master/arabidopsis/summarize_psegs_16jan2018.pl*). Protein domains from Pfam 31 ^66^ were predicted with hmmscan from HMMER 3.1b2 with the argument ‘*--cut_ga*’, which imposed domain-specific thresholds for significance; Pfam domain hits were tabulated with *pfam_hmmscan2annot.pl* (https://github.com/SchwarzEM/ems_perl/blob/master/ *afd2solexa_data/pfam_hmmscan2annot.pl*). Protein domains from InterPro 5.18-57.0 ^67^ were predicted via InterProScan (*interproscan.sh*) with default thresholds and the arguments ‘*-dp -hm - iprlookup -goterms*’. To enable InterProScan searches, which had an upper limit of 1,000 protein sequences per search, the *A. thaliana* proteome was split into 30 equally numerous subsets with *split_fasta_sizewise.pl* (argument ‘*-b 30*’; github) and then searched in parallel; InterPro domain hits were collected and tabulated with *tabulate_iprscan_tsv_22jan2018.pl* (https://github.com/ *SchwarzEM/ems_perl/blob/master/arabidopsis/tabulate_iprscan_tsv_22jan2018.pl*). For both Pfam and InterPro domains, domain predictions were run on the official TAIR10 set of representative (largest) protein isoforms (downloaded from http://www.arabidopsis.org/ *download_files/Sequences/TAIR10_blastsets/TAIR10_pep_20110103_representative_gene_ model_updated*; version dated 4/16/2012). Transcription factor annotations were imported from the AtTFDB and PlantTFDB databases as in ^58^. Gene Ontology (GO) annotations ^68^ of gene function in this column were produced in July 2019 by the *Arabidopsis* genome database, and were downloaded from the GO database for use here. All individual annotations were merged into a single data table with *add_tab_annots.pl* (https://github.com/SchwarzEM/ems_perl/blob/master/ *afd2solexa_data/add_tab_annots.pl*).

Significantly upregulated and downregulated (padj ≤ 0.1) genes in wild-type versus *drmy1-2* mutant inflorescences were separately analyzed with AgriGO (http://systemsbiology.cau.edu.cn/ *agriGOv2*) ^69^ to identify significantly enriched GO terms. Selected GO terms were graphed.

#### Quantification of DR5 and TCS signals

In order to quantify the DR5 and TCS signals at the different positions relative to the center of the floral meristem, we manually aligned the stacks in MorphoGraphX. They were placed with the z-axis located at the meristem center pointing upwards and the x-axis representing the 0° position pointing to the right. Angles increased in the counter-clockwise direction within the xy-plane. The images were aligned in such a way as to place outer sepal position at roughly 45° (the top-right direction when viewing down the z-axis).

For the quantification of the signal, we implemented a custom process in MorphoGraphX which computed a circular histogram of the signal sum around the z-axis. For each voxel of the aligned image, its angle about the z axis was determined. The voxels were grouped according to the angular values in bins of the size of 1° and their signal values weighted by their volume were summed up for each bin. To create the plot, we exported the resulting histogram to a csv-file and imported it into Microsoft Excel.

#### Accession Numbers

RNAseq data: NCBI BioProject PRJNA564625. Individual RNA-seq read sets are archived in SRA as follows: WT replicate 1, SRX6821462, WT replicate 2, SRX6821463, WT replicate 3, SRX6821464, *drmy1-2* replicate 1, SRX6821465, *drmy1-2* replicate 2, and SRX6821466, *drmy1-2* replicate 3, SRX6821467.

Gene names: DRMY1, AT1G58220; AHP6, AT1G80100; TIR1, AT3G62980; AFB1, AT4G03190; AFB2, AT3G26810; AFB3, AT1G12820; WOL, AT2G01830; and PIN1, AT1G73590.

---

**Movie S1.** 6-hour live imaging of one WT flower expressing a plasma membrane marker (*35S::mCitrine-RCI2A*, *pLH13*, white) to track the sepal initiation events. Scale bars: 20 µm.

**Movie S2.** 6-hour live imaging of one *drmy1-2* flower expressing a plasma membrane marker (*35S::mCitrine-RCI2A*, *pLH13*, white) to track the sepal initiation events. Scale bars: 20 µm.

**Movie S3.** 6-hour live imaging of one WT flower with both DR5 (Red) and TCS (Green) to track the spatiotemporal distribution of auxin and cytokinin responses. Purple: chlorophyll auto-fluoresce for flower morphology; Scale bars: 10 µm.

**Movie S4.** 6-hour live imaging of one WT flower with DR5 (White) and plasma membrane marker (*p35S::mCherry-RCI2A*, Red) to track the spatiotemporal distribution of auxin responses. Scale bars: 10 µm.

**Movie S5.** 6-hour live imaging of one WT flower with TCS (green) to track the spatiotemporal distribution of cytokinin responses. Red: chlorophyll auto-fluoresce for flower morphology; Scale bars: 10 µm.

**Movie S6.** 6-hour live imaging of one *drmy1-2* flower with DR5 (White) and plasma membrane marker (*p35S::mCherry-RCI2A*, Red) to track the spatiotemporal distribution of auxin responses. Scale bars: 10 µm.

**Movie S7.** 6-hour live imaging of one *drmy1-2* flower with TCS (green) to track the spatiotemporal distribution of cytokinin responses. Red: chlorophyll auto fluoresce for flower morphology; Scale bars: 20 µm.

**Data S1.** Gene annotations and RNA-seq data for wild-type inflorescences versus *drmy1-2* inflorescences.

This data file contains two tables. The first (n“RNA-seq annots”) enumerates all protein-coding and control genes analyzed by RNA-seq, with identification numbers, aliases, descriptions, motifs, RNA-seq expression data, and instances of significantly changed gene expression between genotypes or batches. The second (“Read counts and mapping”) gives read counts, read lengths, total sequence amounts, and mapping rates for each RNA-seq replicate. Data columns in the annotation table are as follows:

**Gene:** Generally, a given protein-coding gene in the TAIR10 release of the *Arabidopsis* genome database. All further data columns are pertinent to that particular gene; in particular, RNA-seq expression values for each gene were computed with Salmon. In addition to protein-coding genes from *Arabidopsis*, a coding sequence for GFP was also included in this gene set and had its RNA-seq expression values computed as a negative control for background noise (given that GFP was known to be a true negative in all expression sets). Gene names follow standard genome-based identifications (e.g., “AT1G58220” rather than “DRMY1”).

**Primary_name:** The official, primary, human-readable name for a given gene (if any has been designated; e.g., “DRMY1” rather than “AT1G58220”).

**All_names:** The full list of human-readable names for a given gene; this includes not only the primary name, but any aliases in the literature.

**Description:** Short descriptions of gene function, annotated in TAIR.

**Prot_size:** This shows the full range of sizes for all protein products from a gene’s predicted isoforms.

**Max_prot_size:** The size of the largest predicted protein product.

**Phobius:** This denotes predictions of signal and transmembrane sequences made with Phobius 1.01 ^63^. ‘SigP’ indicates a predicted signal sequence, and ‘TM’ indicates one or more transmembrane-spanning helices, with N helices indicated with ‘(Nx)’. Varying predictions from different isoforms are listed.

**Ncoils:** This shows coiled-coil domains, predicted by ncoils ^64^. Both the proportion of such sequence (ranging from 0.01 to 1.00) and the exact ratio of coiled residues to total residues are given. Proteins with no predicted coiled residues are blank.

**Psegs:** This shows what fraction of a protein is low-complexity sequence, as detected by pseg ^65^ As with Ncoils, the relative and absolute fractions of each protein’s low-complexity residues are shown.

**Pfam:** Predicted protein domains from Pfam 31 ^66^, with a domain-specific threshold for significance.

**InterPro:** Domains from InterPro 5.18-57.0 ^67^, predicted with InterProScan using default settings.

**TF:** Annotations in this column denote whether a gene is predicted to encode a transcription factor, as defined in either of two databases for *Arabidopsis* (AtTFDB and PlantTFDB); the particular subtype of transcription factor is noted in brackets. For instance, ATML1 is annotated with “TF [HD-ZIP; Homeobox]”.

**GO_term:** Gene Ontology (GO) annotations ^68^ of gene function in this column were produced in July 2019 by the *Arabidopsis* genome database, and were downloaded from the GO database for use here.

**[*RNA-seq read set*]_TPM:** For the gene in question, and for a given RNA-seq data set, this denotes the gene’s activity as measured in that RNA-seq data set and computed by Salmon in Transcripts Per Million (TPM). RNA-seq data sets were generated for the genotypes WT, and *drmy1-2* (*vos2*); for each genotype, there were three batch replicates (denoted by the suffixes ‘_rep1’ through ‘_rep3’).

**WT.vs.drmy1-2.log2FoldChange:** For a given gene, this denotes the fold change of gene activity, computed by DESeq2 for significant differences between the conditions WT vs. *drmy1-2* (*vos2*) DESeq2 normalized read counts between the three batches for each condition before computing an overall log2FoldChange between two conditions. In each case, a positive value indicates that the first condition has stronger expression than the second condition.

**WT.vs.drmy1-2.pvalue:** For a given gene, this denotes the uncorrected p-value for any significant change of gene activity between two conditions, as listed above for “log2FoldChange[*condition*]”.

**WT.vs.drmy1-2.padj:** For a given gene, this denotes the p-value (adjusted for multiple hypothesis testing, via the Benjamini-Hochberg correction; hence, abbreviated as “padj”) for any significant change of gene activity between two conditions, as listed above for “log2FoldChange[*condition*]”.

**[*RNA-seq read set*]_reads:** For the gene in question, and for a given RNA-seq data set, this denotes the number of RNA-seq reads mapped to the gene (rounded to the nearest integer). These read counts were used to compute fold changes of gene activity between WT and *drmy1-2* genotypes, as well as to compute the significances (p-value and adjusted p-value) of such changes.

**Fig. S1.**
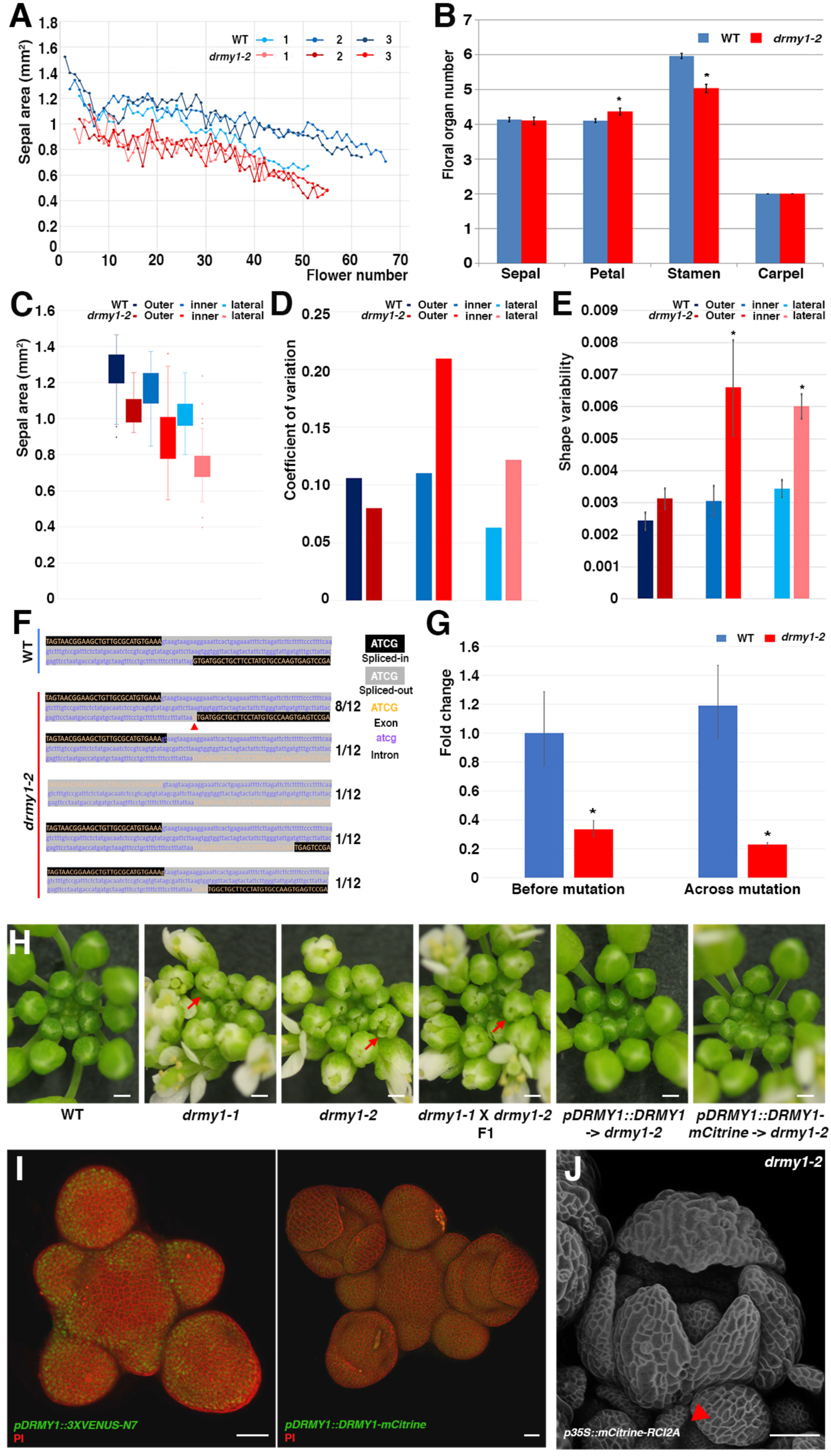
D*R*MY1 is required for sepal size and shape robustness. (A) Quantification of the mean sepal area of the four sepals from an individual flower. Sequential flowers along the main branch of the stem (flower number on the x-axis) were measured at stage 14. Three replicates are included for both WT and *drmy1-2*. (B) *DRMY1* mutations have little effect on the floral organ number. Numbers of sepals, petals, stamens and carpels were quantified for both WT and the *drmy1-2* mutant. Student’s t test * p-value < 0.05. Error bars: standard error of the mean. (C) The sepal area distribution for outer, inner and lateral sepals. The boxes extend from the lower to upper quartile values of the data and the whiskers extend past 1.5 of the interquartile range. Small dots for each box indicate the outliers. Sepals from different flowers were pooled together. n = 48 for both WT and *drmy1-2* 10^th^ to 25^th^ flowers along the main branch. (D) Coefficient of variation (CV) calculated for the areas of the outer, inner and lateral sepals. Sepals from different flowers were pooled together. n = 48 flowers. (E) Quantification of the sepal shape variability for outer, inner and lateral sepals. Student’s t test * p-value < 0.05. Error bars: standard error of the mean. n = 60 for both WT and *drmy1-2* 10^th^ to 25^th^ flowers along the main branch. (F) Sequencing of *DRMY1* transcripts from the *drmy1-2* mutant verified splicing defects occur. *DRMY1* transcripts were reverse transcribed and amplified from RNA extracted from the *drmy1-2* mutant and inserted into *pENTR-D-TOPO* for sequencing. Black shading: nucleotides remaining in the transcript after the splicing; Gray shading: nucleotides spliced out; Orange capital letter: exon; Purple lower-case letter: intron in the WT *DRMY1* transcript. Red arrowhead indicates one base pair shift. (G) qRT-PCR measuring the expression of *DRMY1* in WT and the *drmy1-2* mutant using two pairs of primers: one before the mutation site and another across the mutation site. The expression level in WT quantified with the primers before the mutation site was set to 1 using the Delta-delta-CT method. Student’s t test * p-value < 0.05. Error bars: standard error of the mean. n = 3 biological replicates. (H) Inflorescences of WT, *drmy1-1*, *drmy1-2*, F1 of the cross between *drmy1-1* and *drmy1-2* for allelism test, T3 plants of *drmy1-2* transformed with *pDRMY1::DRMY1*, and T3 plants of *drmy1-2* transformed with *pDRMY1::DRMY1-mCitrine*. Red arrows: smaller sepals in individual flowers. Note, open flower buds indicate unequal sepal sizes. Scale bars: 0.5 mm. (I) Transcriptional (*pDRMY1::3XVENUS-N7*, nuclear localized green signal) and translational (*pDRMY1::DRMY1-mCitrine*, green) *DRMY1* reporter expression patterns are similar. Cell walls were stained with PI. Both *DRMY1* reporters are expressed in the inflorescence meristem, floral meristems, and initiating floral organs, with stronger expression in the periphery. Scale bars: 20µm. (J) What at first appeared to be two sepals initiated at the inner sepal position of the *drmy1-2* flower fused to form a single sepal with a split tip at later time points of the live imaging. Red arrowheads: the fused sepal; Scale bar: 50 µm.

**Fig. S2.**
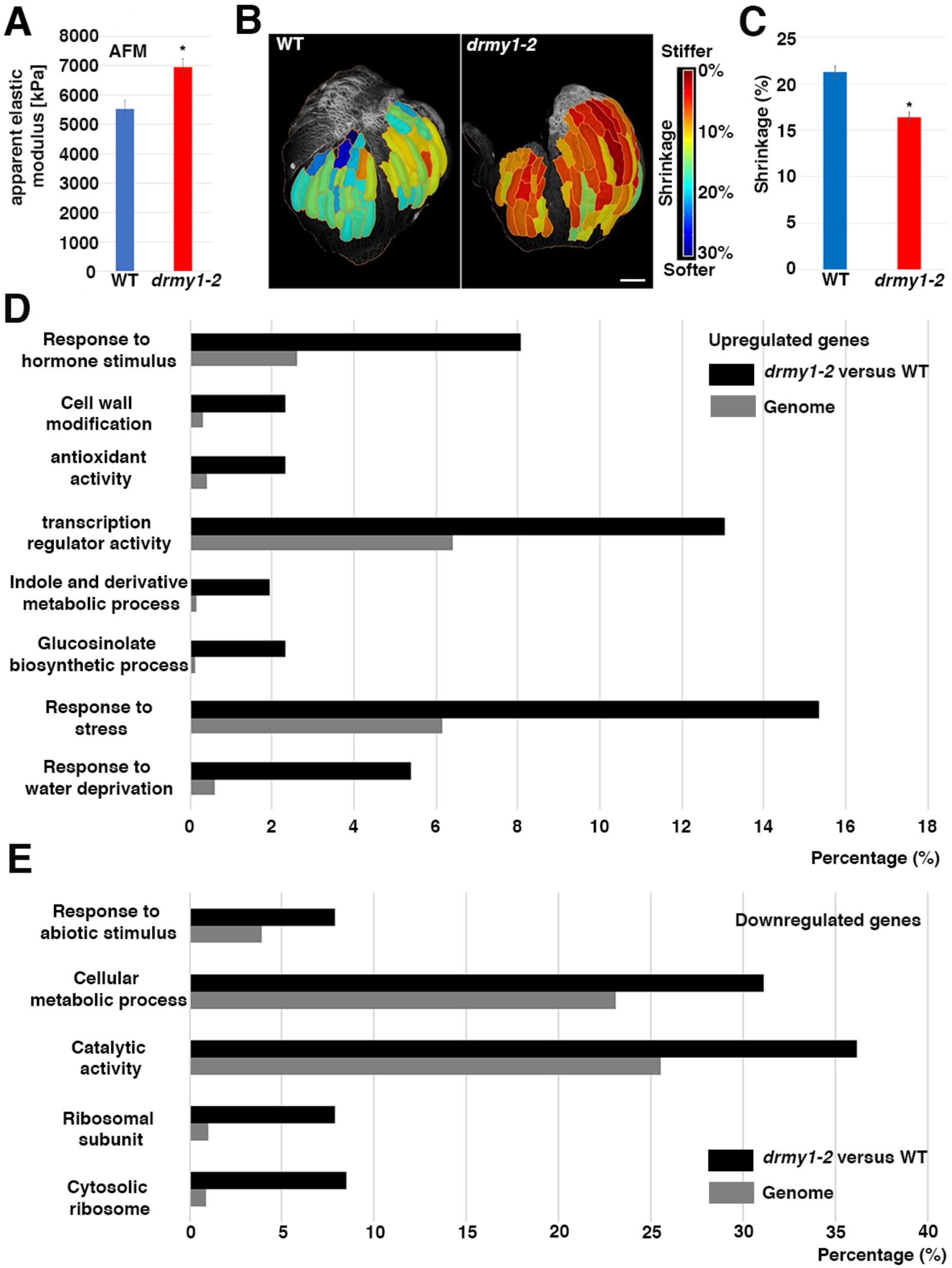
Cell wall stiffness increases in the *drmy1-2* mutant. (A) The average apparent elastic modulus calculated from AFM measurements of the inflorescence meristem, flower meristem, and initiating sepal is significantly higher for the *drmy1-2* mutant. Student’s t test * p-value < 0.05. Error bars: standard errors of the mean. n = 11 samples measured for both WT and *drmy1-2*. (B) Cell shrinkage heatmap after osmotic treatment in WT and *drmy1-2*. Group of cells were segmented together for area comparison. Red in the heatmap represents less shrinkage, thus stiffer cell wall. Scale bar: 50 µm. (C) Average shrinkage ratio after osmotic treatment further confirmed cells undergo less shrinkage in the *drmy1-2* mutant, indicating the cell walls are stiffer. Student’s t test * p-value < 0.05. Error bars: standard errors of the mean. (D) Bar graph of GO terms that are overrepresented (against a genome wide average) among genes more strongly expressed in *drmy1-2* inflorescences than in WT inflorescences. (E) Bar graph of GO terms that are overrepresented (against a genome wide average) among genes more strongly expressed in WT inflorescences than in *drmy1-2* inflorescences For both (D) and (E), a subset of significant GO terms was selected for each graph. Genome (gray) reports the frequency of genes associated with that term in the *Arabidopsis* genome, which would be the frequency expected by chance for a randomly selected subset of genes. The percentage of genes was calculated as the number of genes associated with that term divided by the total number of genes

**Fig. S3.**
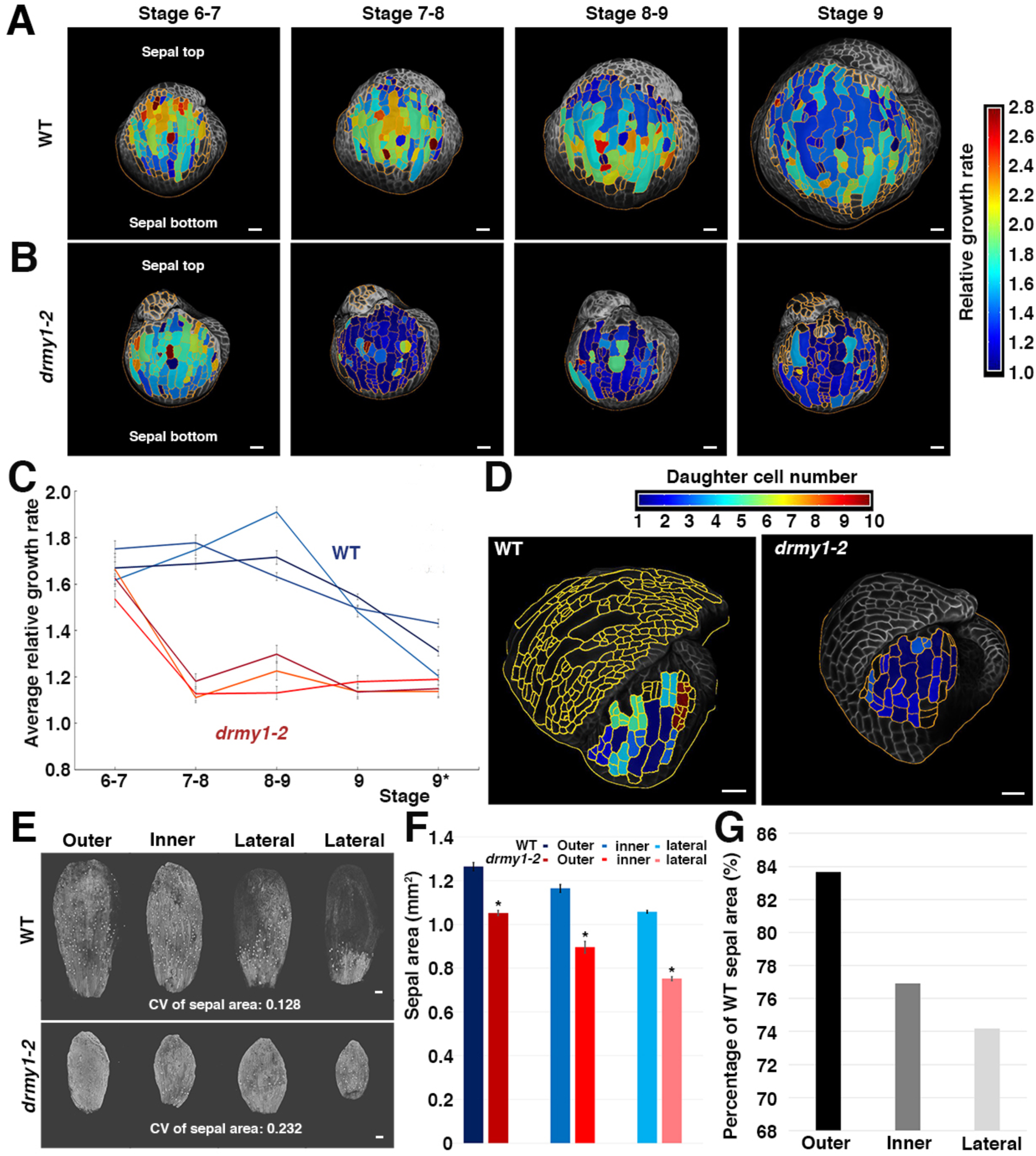
Sepal cell growth is slower in the *drmy1-2* mutant. (A and B) 24-hour late stage (from stage 6 to stage 9) cellular growth heatmap for both WT (A) and *drmy1-2* (B) outer sepals. Relative growth rate is defined as final cell size divided by initial cell size. Segmented cells outlined in yellow. Note the outer sepal base is at the bottom of the image and its tip points up. Scale bars: 20 µm. (C) Growth curves of the late stage average cellular growth for both WT and *drmy1-2*. *: Flower stage 9 extends over multiple 24-hour intervals. Error bars represent standard error of the mean. (D) 36-hour cell division heatmap for both WT and *drmy1-2*. The total number of cells derived from one progenitor is represented in the heatmap with 1 meaning no divisions. Throughout sepal development, the *drmy1-2* sepal cells undergo fewer divisions than WT. Scale bars: 20 µm. (E) Confocal images of sepals from individual flowers (shown in Fig. 3A-B) after 11 days of live imaging. Area variability was quantified by the coefficient of variation (CV). The first *drmy1-2* lateral sepal fused from two lateral primordia. Scale bar: 100 µm. (F) The mean sepal area for WT and *drmy1-2* outer, inner and lateral sepals. Sepals from different flowers were pooled together. n = 48 flowers. Student’s t test * p-value < 0.05. Error bars represent standard error of the mean. (G) The mean *drmy1-2* sepal area divided by WT mean sepal area ratio for each sepal type. Sepals from different flowers were pooled together. n = 48 flowers.

**Fig. S4.**
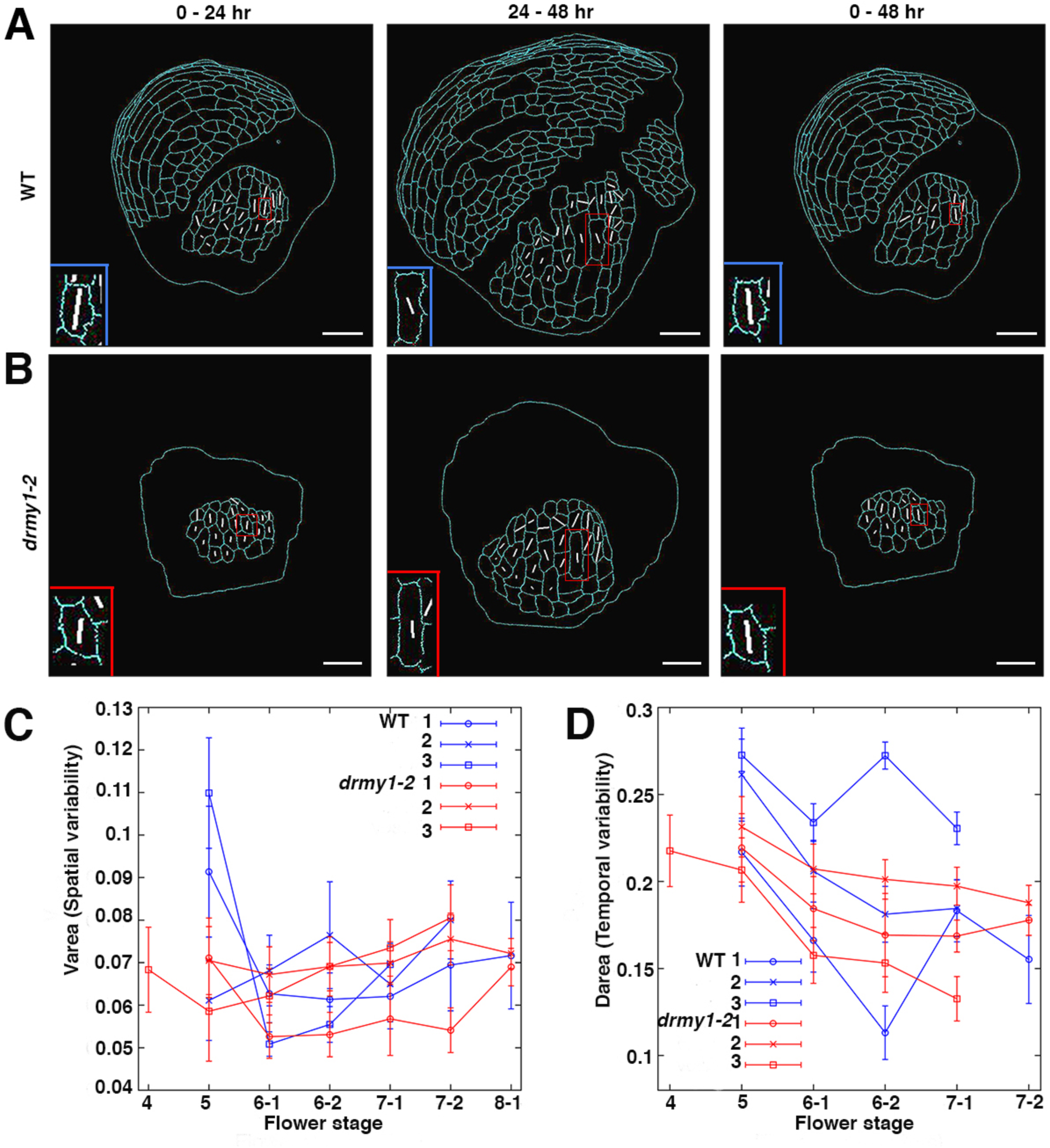
Spatiotemporal averaging is not affected in the *drmy1-2* mutant at early stage. (A) The maximal principal direction of growth (PDG_max_, white line) of WT sepal cells calculated for 24-hour and 48-hour intervals. For 24-hour intervals, the PDG_max_ shows both spatial and temporal variations in WT. Cell outlines are shown in cyan. Over the whole 48-hour interval these variations average out such that the PDG_max_ are oriented vertically along the major growth axis of the sepal. One cell showing nice temporal averaging is highlighted with blue boxes and magnified in insets. Scale bar: 20 µm. (B) The maximal principal direction of growth (PDG_max_, white line) of the *drmy1-2* sepal cells calculated for 24-hour and 48-hour intervals. The PDG_max_ also shows similar spatial and temporal variations in the *drmy1-2* situation. One cell showing nice temporal averaging is again highlighted with red boxes at different time points, indicating the temporal averaging of growth direction is not affected by *DMRY1* mutations and magnified insets. Scale bar: 20 µm. (C) Graph plotting the average spatial variability of the growth rates (V_area_) for sepal epidermal cells during the development of sepals. Blue curves are for WT sepals and red curves are for the *drmy1-2* sepals. (D) Graph plotting the average temporal variability of the growth rates (D_area_) for sepal epidermal cells during the development of sepals. Blue curves are for WT sepals and red curves are for the *drmy1-2* sepals.

**Fig. S5.**
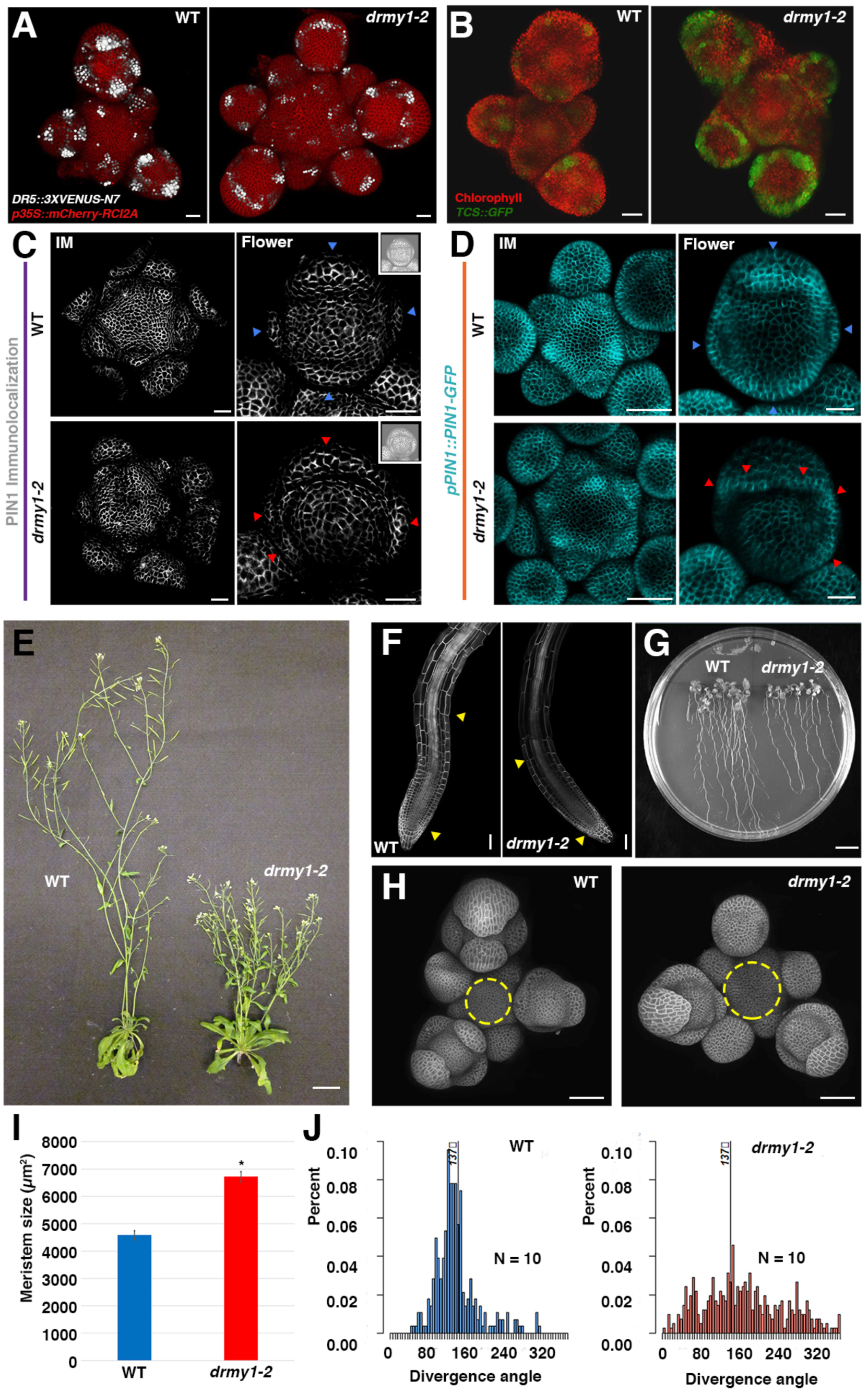
Auxin signaling is suppressed and more diffuse while cytokinin signaling expands and is enhanced in *drmy1-2* mutants. (A) Confocal imaging of the DR5 auxin response reporter (white) in the whole inflorescence of WT and the *drmy1-2* mutant. *p35S::mCherry-RCI2A*: red, for plasma membrane. Scale bar: 20µm. (B) Confocal imaging of the TCS cytokinin signaling reporter (green) in whole inflorescences of WT and the *drmy1-2* mutant. Chlorophyll autofluorescence: red. Scale bar: 20 µm. (C) Confocal imaging of PIN1 immunolocalization experiments to show PIN1 accumulation in inflorescences and flowers of WT and *drmy1-2*. PIN1 exhibits polar localization in *drmy1-2* similar to wild type; however, it forms more convergence points in flowers. Blue/Red arrowheads: PIN1 convergence points. Inset: Same images in B and F with increased brightness to show the structure of the flowers; Scale bars: 20 µm. (D) Confocal imaging of *pPIN1::PIN1-GFP* to show PIN1 accumulation in the inflorescences and flowers of WT and *drmy1-2*. Again, PIN1 form abnormal convergence points in *drmy1-2*. Blue/Red arrowheads: PIN1 convergence points; Scale bars: 20 µm. (E) Images of whole plants for WT and *drmy1-2,* showing the bushiness and short stature of *drmy1-2*. Scale bar: 2 cm. (F) Confocal images of root meristems for WT and *drmy1-2*. The regions specified by yellow arrowheads indicate the meristematic zone. Scale bar: 50 µm. (G) Photograph of 10-day old seedlings for WT and *drmy1-2,* showing *drmy1-2* has shorter roots and fewer lateral roots. Scale bar: 1 cm. (H) Confocal images of inflorescence meristems for WT and *drmy1-2* (Top view). Yellow dashed circles indicate how meristem sizes were measured in I. Scale bar: 50 µm. (I) Quantification of inflorescence meristem sizes for WT and *drmy1-2*. Student’s t test * p-value < 0.05. (J) Histograms of divergence angles between siliques for WT and *drmy1-2*, showing the enhanced variability in phyllotaxy observed in *drmy1-2* mutants. 137° is expected for spiral phyllotaxy observed in wild type.

**Fig. S6.**
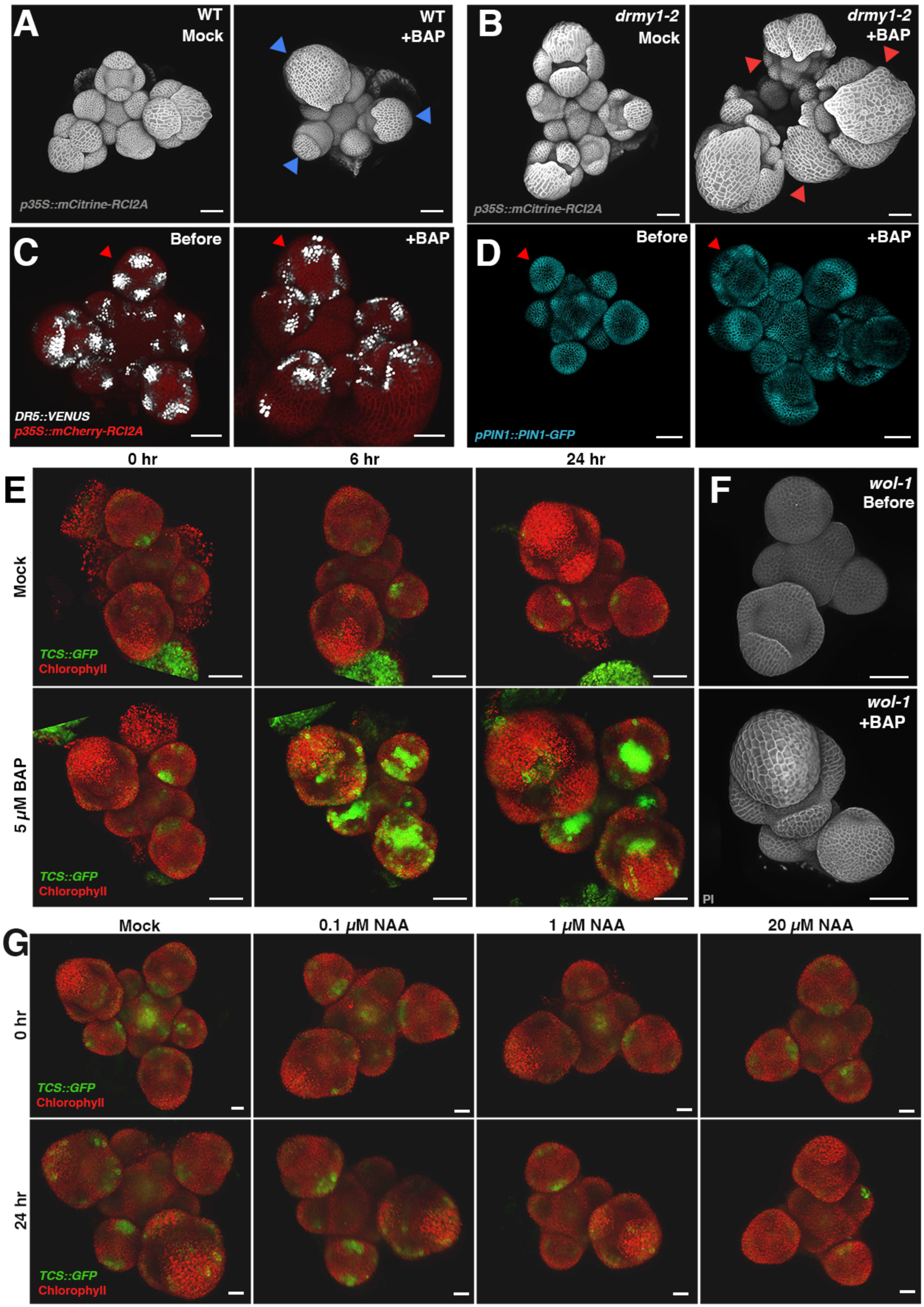
Cytokinin treatment mimics the *drmy1-2* mutant. (A and B) Confocal imaging of the whole inflorescences of WT (A) and *drmy1-2* (B) cultured in mock conditions or 5 µM BAP (synthetic cytokinin) for 6 days. *p35S::mCitrine-RCI2A*: gray, for plasma membrane. Blue or red arrowheads: flowers with obvious delayed sepal initiation phenotypes. Scale bars: 50 µm. (C) 5 µM BAP treatment on the DR5 auxin signaling reporter (white) for 3 days. *p35S::mCherry-RCI2A*: red, for plasma membrane; Red arrowhead: indicates the same flower before and after the BAP treatment. Scale bar: 50 µm. Note the DR5 signal becomes more diffuse after cytokinin treatment. (D) 5 µM BAP treatment on PIN1-GFP (cyan) auxin efflux carrier for 2 days. Red arrowhead: indicates the same flower before and after the BAP treatment; Scale bar: 50 µm. PIN1-GFP appears to form additional convergence points similar to *drmy1-2*. (E) 5 µM BAP treatment on the TCS cytokinin signaling reporter (green) for 24 hours. Control showing that cytokinin treatment enhances TCS reporter expression. Chlorophyll autofluorescence: red; Scale bars: 50 µm. (F) 5 µM BAP treatment on the cytokinin receptor mutant *wol-1* for 4 days. Control showing that mutation of the cytokinin receptor (*wol-1*) abrogates delayed sepal initiation in response to cytokinin. Lower left flower removed during imaging. Cell walls stained with PI: gray; Scale bar: 50 µm. (G) NAA (auxin) treatment in a gradient of concentration on the TCS cytokinin signaling reporter (green) for 24 hours. Auxin treatment did not enhance TCS reporter expression. Chlorophyll autofluorescence: red; Scale bars: 20 µm.

**Fig. S7.**
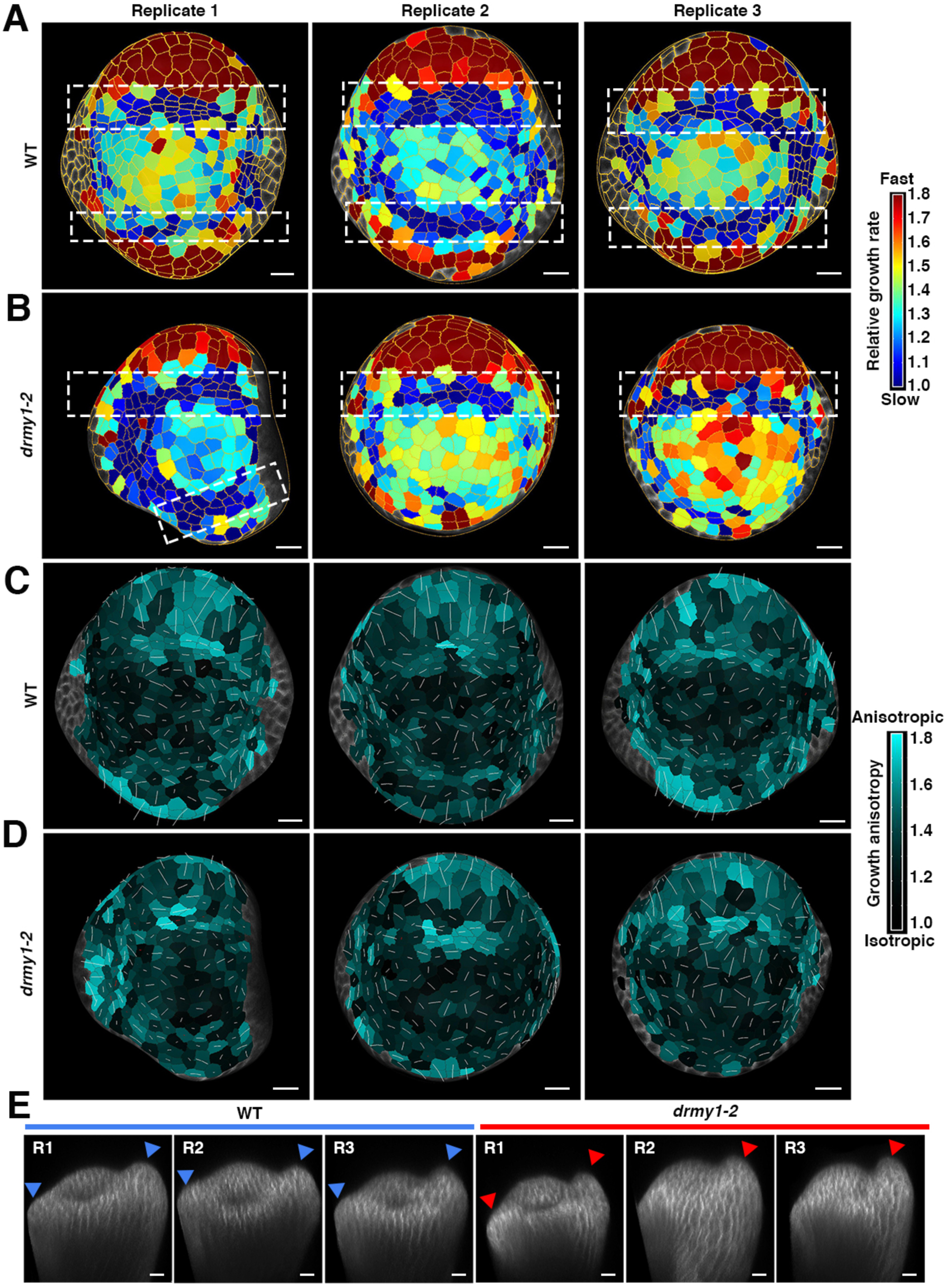
Cellular growth remains heterogeneous and randomly oriented for the *drmy1-2* inner sepals. (A and B) Cumulative 18-hour cellular growth heatmap for both WT (A) and *drmy1-2* (B) floral meristems. White dashed boxes highlight the bands of cells with slower growth rate which specify the boundary. They are always adjacent to the fast growth regions at the periphery, where sepals initiate. Segmented cells outlined in yellow. Three replicates are shown. Note that the *drmy1-2* replicate 1 grows relatively normally. Scale bars: 10 µm. (C and D) 18-hour cellular growth anisotropy heatmap for the same WT (C) and *drmy1-2* (D) floral meristems. Growth anisotropy was calculated by dividing the cell stretch at the maximum direction by the cell stretch at the minimum direction. Cyan indicates higher growth anisotropy while black indicates lower growth anisotropy. White lines within the cells shows the maximum principle directions of growth. The initiating regions have higher anisotropy with the periphery part showing longitudinal growth and the boundary parts showing latitudinal growth. Scale bars: 10 µm. (E) Side views of the floral meristems at the last time points with outer sepals on the right and inner sepals on the left. The morphology of outer sepals was used for staging and appears equivalent in all samples. Arrowheads: the initiation/bulging of sepals from floral meristems. Scale bars: 10 µm.

**Table S1.**
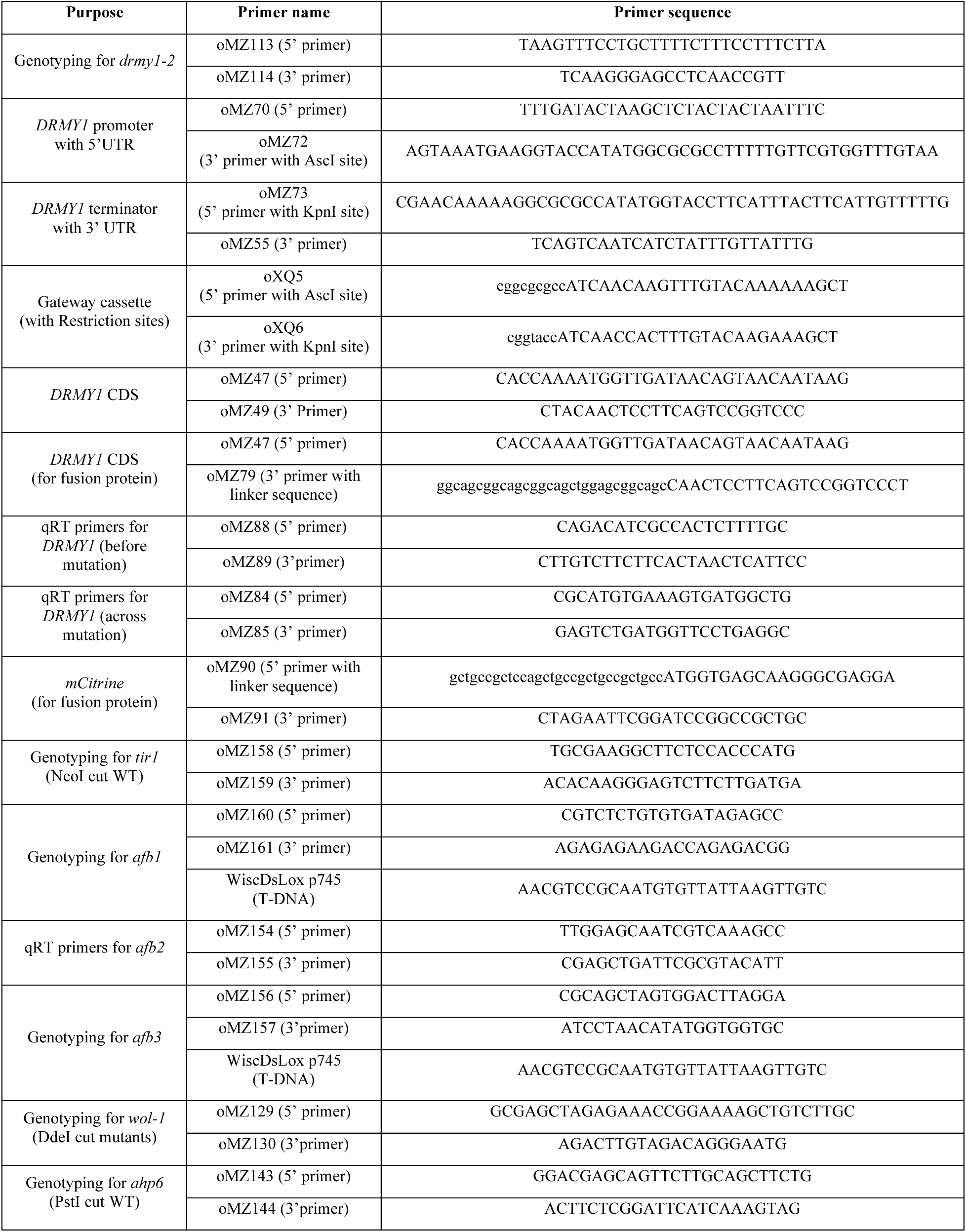
Information about the primers used in this paper.

**Table S2.** Information about the fluorescent signals used in this paper.

| Fluorescent signals | Excitation | Emission |
| --- | --- | --- |
| mCitrine/VENUS | 514 nm | 519 - 621 nm |
| VENUS (when together with mCherry) | 514 nm | 519 - 578 nm |
| mCherry | 594 nm | 599 - 687 nm |
| GFP | 488 nm | 493 - 598 nm |
| GFP (when imaged together with VENUS) | 488 nm | 493 - 523 nm |
| Chlorophyll | 488 nm | 647 - 721 nm |
| PI | 514 nm | 566 - 656 nm |

**Table S3.**
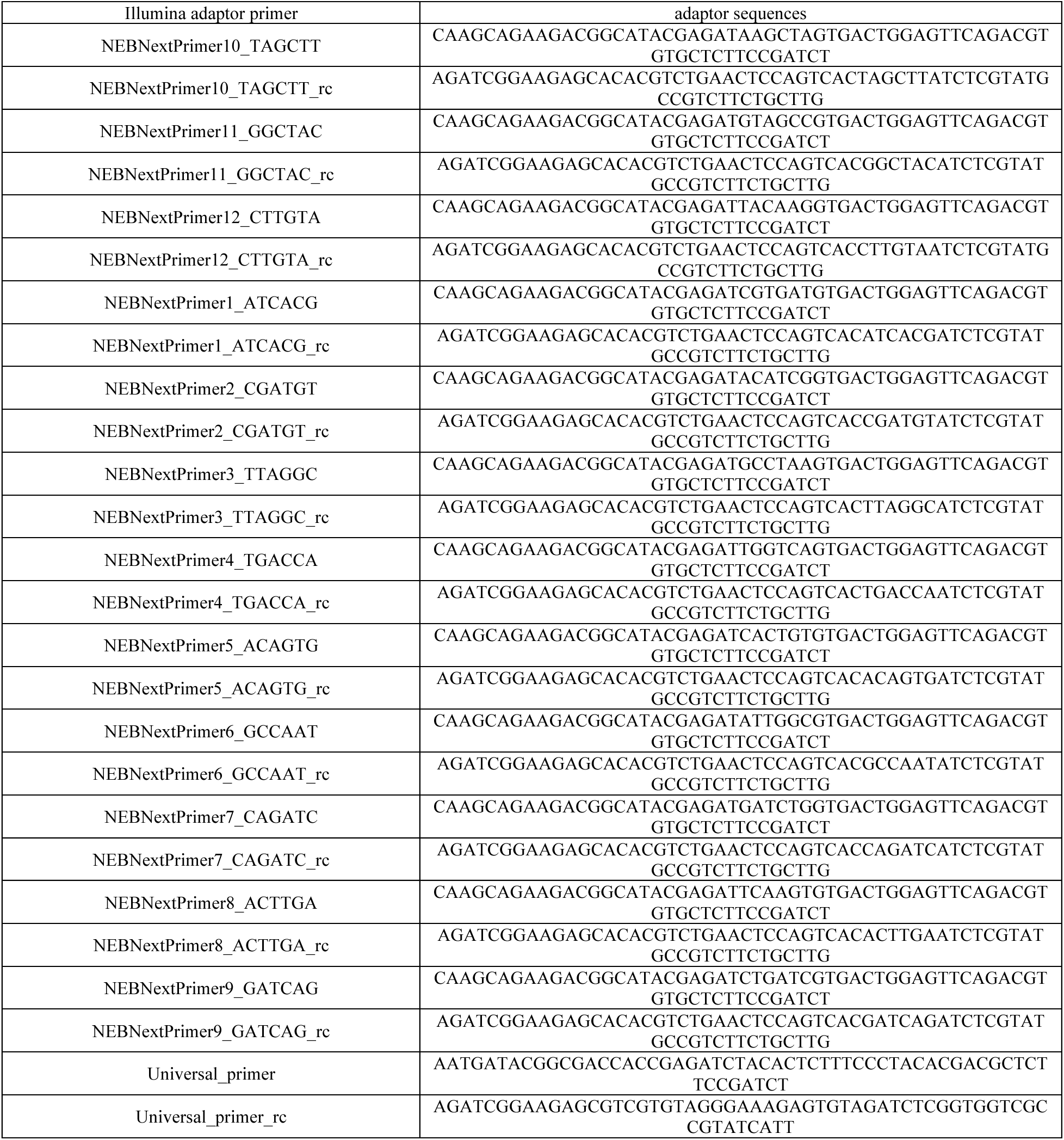
Illumina adaptor sequences (*sepal_adapters_09dec2016.trimmomatic.fa*) used for trimming RNA-seq reads via Trimmomatic.

